# Small Area Estimation of Forest Volume Using Mixed Effects Random Forests and Multi-Source Remote Sensing Data

**DOI:** 10.64898/2026.04.22.720077

**Authors:** Elia Vangi

## Abstract

Accurate estimation of forest growing stock volume (GSV) at fine spatial scales is essential for sustainable forest management, carbon accounting, and local decision-making. However, traditional forest inventories often lack sufficient sampling density to provide reliable estimates for small areas. This study evaluates the performance of two small area estimation approaches: the Empirical Best Predictor (EBP) based on a nested-error linear regression model, and the Mixed-Effects Random Forest (MERF) for estimating GSV at the forest stand level using multi-source remote sensing data. The analysis was conducted in the Vallombrosa Nature Reserve (Italy), integrating field measurements from 101 plots with auxiliary variables derived from Sentinel-2 imagery and airborne LiDAR. Both methods were applied to estimate the mean and total GSV across 658 forest stands, many of which lacked direct observations. Model performance was assessed using spatial cross-validation, and uncertainty was quantified using root-mean-square error (RMSE). Results show that MERF outperformed EBP in predictive accuracy, achieving higher R^2^ (0.67 vs. 0.37) and lower RMSE (151 vs. 202 m^3^ ha□^1^). MERF also produced more stable and precise uncertainty estimates, with improved coverage of observed values. While both methods yielded comparable total GSV estimates, EBP exhibited greater variability and sensitivity to model assumptions. In contrast, MERF effectively captured non-linear relationships and handled multicollinearity among predictors, though at the cost of reduced interpretability and higher computational demand. Overall, findings highlight the advantages of integrating machine learning with mixed-effects modeling for SAE in forestry, particularly under conditions of sparse sampling and complex ecological variability.

## 1. Introduction

Forests are a fundamental element of climate change mitigation and biodiversity conservation efforts, and monitoring them is crucial for promoting sustainable management, carbon accounting, and ecosystem service evaluation. Traditional national forest inventories (NFIs), such as those implemented in many European countries, typically rely on sparse ground plot networks and design-based inference techniques (Tomppo et al., 2010; McRoberts et al., 2009). While these approaches enable unbiased estimates of forest parameters at national or regional levels, they are inadequate for generating reliable data at smaller administrative or ecological units, such as municipalities or protected areas, due to limited plot density.

As decision-making processes become more decentralized and spatially detailed, there is an increasing need for forest information at local levels, along with reliable uncertainty estimates. This need arises at two related levels. On the one hand, countries require robust estimates of forest variables (e.g., growing stock volume, carbon stocks) for their international reporting obligations under frameworks such as the UNFCCC, LULUCF, and REDD+. These estimates must explicitly quantify uncertainty, in line with the IPCC’s good-practice guidelines. On the other hand, subnational actors, such as municipalities, regions, and forest managers, require reliable, detailed spatial data to support operational decisions and policymaking. Local estimates and maps of forest resources are essential for several forest activities, including management, conservation, risk assessment, and quantifying ecosystem services. Local forest management plays a crucial role in sustainable land-use planning, as it provides the foundation for balancing ecological, economic, and social objectives at the landscape scale. Accurate knowledge of growing stock volume (GSV) and above-ground biomass (AGB) is fundamental in this context, as it directly informs decisions on harvesting intensity, timing, and the spatial allocation of interventions. By quantifying the available forest resources, forest managers can ensure that harvest levels remain within sustainable limits, avoiding overexploitation while optimizing economic returns. Moreover, reliable GSV estimates support the design of silvicultural treatments, the planning of infrastructure such as forest roads, and the assessment of carbon stocks and ecosystem services. Here, aggregated statistics alone are insufficient; spatially explicit, fine-scale data, preferably linked to measures of reliability and confidence, should be the standard. Addressing both needs simultaneously requires estimation frameworks that combine the statistical rigor of forest inventories with the spatial detail and coverage of remote sensing data.

To address the limitations of traditional inventory methods, the concept of Enhanced Forest Inventories (EFI) has been developed, combining ground measurements with comprehensive remote-sensing data using model-based techniques (White et al., 2016; White et al., 2025). EFI methods have shown their ability to produce detailed forest attribute maps, such as GSV or AGB, by utilizing the predictive capabilities of satellite and airborne data (e.g., Landsat, GEDI, LiDAR), often employing regression or machine learning algorithms (Chirici et al., 2020; Vangi et al., 2023; Saarela et al., 2015).

However, a key challenge remains: while predictive maps provide detailed spatial information, they rarely include reliable uncertainty estimates at aggregated levels (Saarela et al., 2020). Simply aggregating pixel-based predictions across small areas does not yield valid estimates, as it overlooks spatial model bias and fails to account for sampling error.

Small Area Estimation (SAE) provides a statistical framework to address this problem. Originating from official statistics (Rao, 2015; Rao & Molina, 2015), SAE techniques have been adapted for forest inventory applications to generate unbiased and accurate estimates in domains with few or no field plots (Breidenbach & Astrup, 2012; Goerndt et al., 2013; Magnussen & Breidenbach, 2017). SAE methods can operate at the unit or area level. Traditional SAE methods rely heavily on model-based approaches that “borrow strength” across areas by exploiting auxiliary information. Among these, the Empirical Best Predictor (EBP) under unit-level mixed models—most commonly the nested-error linear regression model (Battese et al., 1988)—has long served as a benchmark due to its solid theoretical foundation, design-consistent properties, and relatively straightforward implementation.

Despite its widespread use, the EBP framework is grounded in the restrictive assumptions of the linear model. In particular, it typically relies on linear relationships between the response variable and auxiliary covariates, and on normally distributed, homoscedastic errors. While these assumptions facilitate analytical tractability and allow for closed-form or simulation-based predictors, they may be overly simplistic in many forestry applications characterized by complex, non-linear interactions, high-dimensional covariates, and heterogeneous processes. As a consequence, model misspecification can lead to biased estimates, reduced efficiency, and poor predictive performance, especially when the underlying relationships deviate substantially from linearity (Molina and Martin, 2018; Rojas-Perilla et al., 2020).

These limitations have motivated the development of more flexible, data-driven approaches that can better capture complex patterns while retaining the core idea of borrowing strength across areas. In the context of estimating and mapping socio-economic indicators (i.e., poverty, income, inequality; Hobza and Morales, 2016), as well as in the health care field (i.e., care-cost opportunity work) and policy settings, the Mixed-Effects Random Forest (MERF) has emerged as a promising alternative (Hajjem et al., 2014; Krenmair & Schmid, 2022), demonstrating superior performance compared to classical SAE methodology, in both unit and area-level frameworks. MERF combines the strengths of machine learning methods—specifically, random forests’ ability to model non-linearities and interactions without explicit specification (Breiman, 2001)—with the hierarchical structure of mixed-effects models, which account for area-level heterogeneity through random effects. By iteratively estimating fixed effects via random forests and random effects via mixed-model techniques, MERF provides a hybrid framework that relaxes parametric assumptions while preserving the capacity to account for hierarchical dependencies. The original MERF method has already been improved to better handle multicollinearity by introducing PCA before MERF and replacing RF with rotation forests (Anand et al., 2024). MERF, as an interesting and relatively new method, raises new methodological and practical questions, including interpretability, uncertainty quantification, and computational cost, which are less problematic in the classical EBP framework.

These approaches are especially practical when auxiliary variables, such as canopy height models (CHM), GEDI RH metrics, or spectral indices, are strongly correlated with the target variable. Recent studies highlight the importance of generating not just estimates but also uncertainty maps, such as pixel-level RMSE or confidence intervals (Saarela et al., 2020; Foody, 2024). This information is crucial for transparent reporting, model validation, and making risk-informed decisions in forestry and land use. Furthermore, although high-quality auxiliary data (such as Landsat, Sentinel-1, and Sentinel-2 composites) are becoming increasingly accessible, they have not yet been fully integrated into design-based SAE at the forest stand level. Few studies have examined SAE applications in the Mediterranean context, and, to our knowledge, none have compared the EBP with the MERF framework in forest inventory.

This paper contributes to the growing literature on SAE by providing a comparative analysis of EBP and MERF with particular emphasis on the trade-offs among model interpretability, flexibility, and predictive performance. By highlighting the conditions under which each approach is most appropriate, we aim to clarify the role of modern machine learning methods within the established SAE paradigm. Furthermore, this study aims to bridge the gap between advanced SAE methodologies and their practical application in forest management. We introduce a reproducible approach to estimate GSV and its associated uncertainty at the stand level using field plots and multi-source remote-sensing-based auxiliary variables.

## 2. Materials and methods

### 2.1 Study area

We conducted our study in the Nature Reserve of Vallombrosa (43.745 E, 11.562 N), located in the Tuscany region, on the north-west side of the Pratomagno Massif (Italy) (Figure 1). The Nature Reserve of Vallombrosa covers 1273 ha and ranges in altitude from 470 to 1447 m a.s.l.

**Figure 1.**
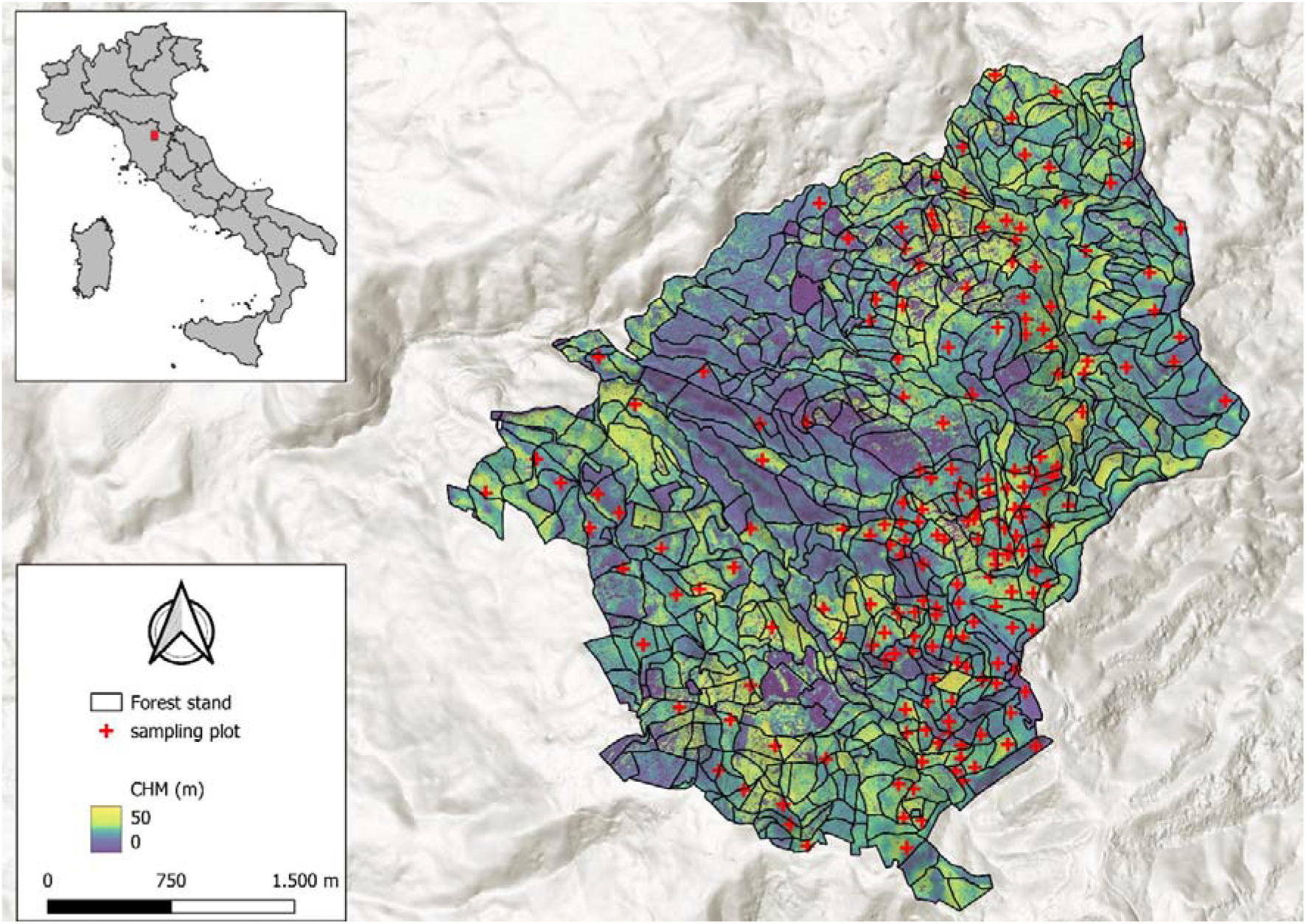
Boundaries of the forest stands and distribution of the sampling field plot. In the background, the CHM.

The dominant forest species are silver fir (Abies alba Mill.) (664 ha), and European beech (Fagus sylvatica L.) stands (187 ha), which mainly originated from coppice conversion to high forest. Traditionally, silver fir has been cultivated in pure stands in the Vallombrosa Forest for many centuries. Other common forest species included *Pinus nigra J*.*F. Arnold, Pseudotsuga menziesii (Mirb*.*) Franco*, introduced for experimental purposes, and *Castanea sativa Mill*., *Quercus cerris L*., *Q. pubescens Willd*., *Ostrya carpinifolia Scop*., *and Fraxinus ornus L*. The forest management plan for the period 2006–2025 (Ciancio, 2009) divided the Reserve into 658 forest stands (i.e., management units) homogeneous in species composition and environmental condition. The mean domain area is 1.9 ha, ranging from 0.15 ha to 13 ha. The Reserve Management plans are currently being revised in preparation for the next management cycle (2026-2046). The next plan will guide the harvesting period, the maximum amount of wood that can be removed from each forest stand, and will set the monitoring system for forest resources and biodiversity. For this reason, detailed spatial information on GSV distribution and its uncertainty will inform plan updating and intervention planning.

### 2.2 Field data

In this study, we used field data from 101 13 m-radius circular sampling plots collected between 2015 and 2016 (Figure 1), which were part of a systematic, unaligned sampling design with a 1 ha hexagonal grid. Within each plot, tree species, diameter at breast height (DBH), tree height, and age were measured and aggregated at the plot level. From structural variables (i.e., DBH and height), growing stock volume (GSV) was computed using species-specific allometric equations developed within the Italian NFI field survey framework (Tabacchi et al., 2011). Only 79 of 658 forest stands (∼12% of the total) have at least one sample plot, with a median of one and a maximum of four plots. In this study, we used GSV plot-level data to estimate the total and mean GSV indicators for all forest stands in the management plan, along with their uncertainty (mean squared error [MSE] and root mean squared error [RMSE]).

### 2.3. Forest mask

To ensure reliable results in small-area GSV estimation, a forest mask was needed to restrict the estimation to forest areas. We used the forest mask produced by D’Amico et al. (2021) and further updated for the reference year 2020 (D’Amico et al., 2023), which closely mimics the standard definition of forest used by the Italian NFI and the FAO, and is also used by the Vallombrosa Reserve management plan.

### 2.4. Remote sensing variables

SAE relies on a statistical model that links field observations to auxiliary variables available at the unit level, i.e., over the entire area of interest (often referred to as wall-to-wall). RS variables meet this fundamental criterion and are usually globally available, offering high spatial and temporal resolution. In this study, a total of 13 RS variables (hereinafter referred to as predictors), mostly derived from the Sentinel-2 mission, were used to fit the nested-error linear regression and random forest models for the EBP and MERF methods, respectively.

#### 2.4.1. Sentinel-2-based predictors

We selected all S2 surface reflectance level 2 images acquired over the study area from the Google Earth Engine archive (Gorelick et al., 2017) from 1^st^ April to 30^th^ September 2015, with less than 50% cloud cover (White et al., 2014). The S2 cloud probability dataset (Shi et al., 2022) was used to mask all pixels with a cloud probability of 65% or higher (Parisi et al., 2023). From each image, six spectral bands were retained (red, green, blue, nir, swir1, and swir2) and five vegetation indices (VIs) were computed (Table 1), resulting in a total of 11bands per image (6 original bands + 5 VIs). We construct a cloud-free summer composite mosaic for each band and VI using the medoid method (Kennedy et al., 2018). The Medoid pixel compositing algorithm seeks to construct the final image using pixels whose surface reflectance values are as close as possible to the median derived from the entire image collection. The distance from the median was calculated via the Euclidean distance.

**Table 1.**
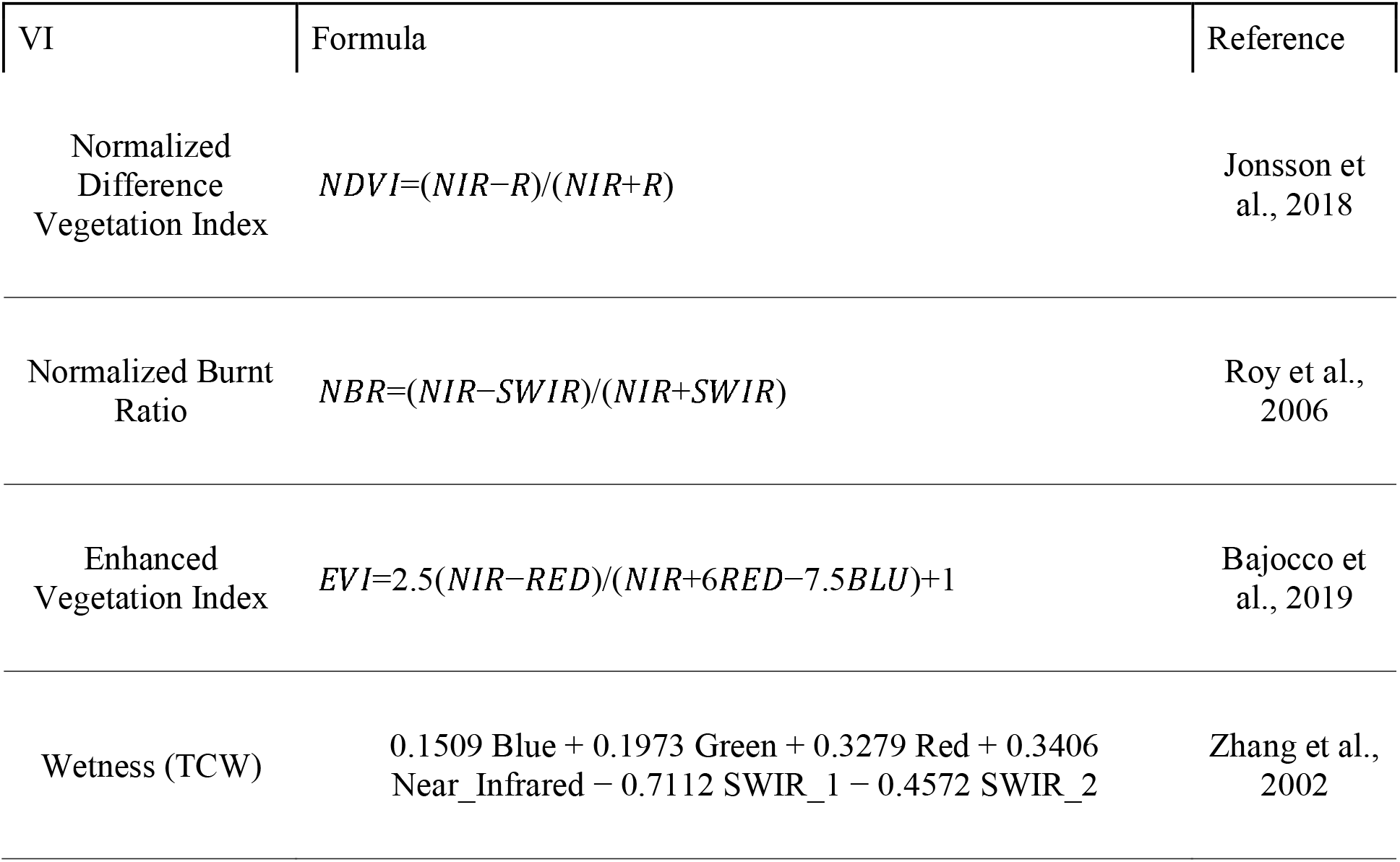
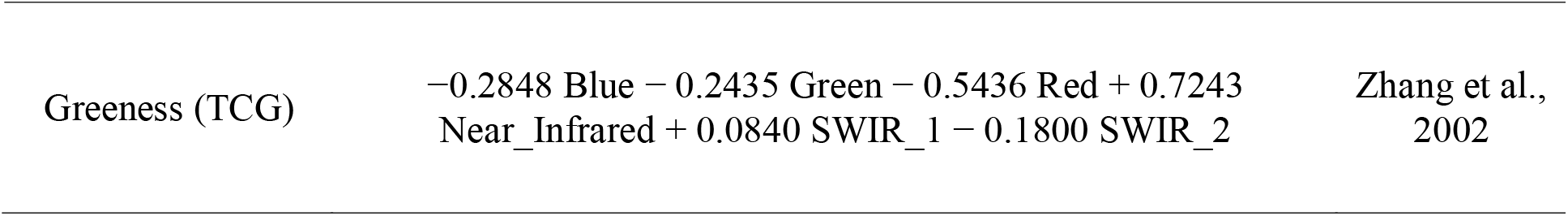
List of vegetation indices used as predictors, along with their formulation and source.

| VI | Formula | Reference |
| --- | --- | --- |
| Normalized<br>Difference<br>Vegetation Index | $NDVI = (NIR - R) / (NIR + R)$ | Jonsson et al., 2018 |
| Normalized Burnt<br>Ratio | $NBR = (NIR - SWIR) / (NIR + SWIR)$ | Roy et al., 2006 |
| Enhanced<br>Vegetation Index | $EVI = 2.5(NIR - RED) / (NIR + 6RED - 7.5BLU) + 1$ | Bajocco et al., 2019 |
| Wetness (TCW) | $0.1509 \text{ Blue} + 0.1973 \text{ Green} + 0.3279 \text{ Red} + 0.3406 \text{ Near\_Infrared} - 0.7112 \text{ SWIR}_1 - 0.4572 \text{ SWIR}_2$ | Zhang et al., 2002 |
| Greenness (TCG) | $-0.2848 \text{ Blue} - 0.2435 \text{ Green} - 0.5436 \text{ Red} + 0.7243$<br>$\text{Near\_Infrared} + 0.0840 \text{ SWIR}_1 - 0.1800 \text{ SWIR}_2$ | Zhang et al.,<br>2002 |

#### 2.4.2. ALS-based predictors

The airborne laser scanning (ALS) dataset was obtained from a survey conducted in May 2015 using a RIEGL LMS-Q680i LiDAR system (Horn, Austria) and a DIGICAM H39 RGB and CIR optical sensor, both installed on an Eurocopter AS350 B3 helicopter. The mission was flown at approximately 1100 m above ground level, at 70 knots, with 30% overlap between adjacent flight lines. The LiDAR system collected full-waveform data with an average point density of 10 points per square meter. Georeferencing of the ALS data was performed in the WGS84 UTM32N coordinate system by refining flight trajectories with Global Navigation Satellite System (GNSS) and Inertial Measurement Unit (IMU) data acquired from two Italtopos network base stations (Giannetti et al., 2018). Standard preprocessing steps—such as outlier and noise removal, classification of ground and non-ground returns, and height normalization—were performed using LAStools (version 240220; Gilching, Germany). Subsequently, digital terrain and digital surface models (DTM and DSM) were generated at a 1 m spatial resolution, and the canopy height model (CHM) was derived by subtracting the DTM from the DSM.

#### 2.4.3. Auxiliary predictors

We complement the set of Sentinel-2+ALS-based predictors by adding the elevation from the DTM and the position (X and Y coordinates). These predictors are strictly related to the position of each unit (i.e., pixel), and incorporating them allows the model to capture part of the spatial structure through the fixed effects component. Although this approach does not fully model spatial dependences, it provides a simple and effective way to mitigate spatial autocorrelation when more complex spatial models are not adopted.

Each predictor was produced at a 10×10 m resolution. Within each sampling plot, we computed the mean values for each predictor in the set.

### 2.5. Methods

#### 2.5.1. The Empirical Best Predictor estimator

The benchmark estimator for the stand-level mean GSV was the EBP, as described by Molina and Rao (2010) and Rao and Molina (2015).

The notation used in this study follows that of Kreutzmann et al. (2019), who developed the R package emdi that implements the EBP. The same notation is used by Krenmair (2022), who developed the R package SAEforest, which implements the MERF algorithm. As demonstrated by Rao and Molina (2015), in the case of linear parameters (i.e., any parameter that can be expressed as a linear combination of unit-level responses), the EBP estimator is equal to the Empirical Best Linear Unbiased Predictor (EBLUP) estimator. Since Battese, Harter, and Fuller (1988), Rao and Molina (2015), and most R packages that implement EBLUP rely on a simplified version in which unit-level auxiliary data are replaced by their domain means (area-level model), we opted for EBP implemented in the emdi package instead, to make full use of the available unit-level auxiliary information.

The finite population of size N (here the total number of pixel in the study area) is denoted by U and partitioned into D domains (the forest stand in this study) U_1_, U_2_, … U_D_ of sizes N_1_, N_2_, … N_D_, where i = 1,…, D refers to an i^th^ domain and j = 1,…, N_j_ to the j^th^ pixel in a domain. From this population, a random sample of size n is drawn. This leads to n_1_,…, n_D_ observations in each domain. If n_i_=0, the i^th^ domain is not in the sample. The target variable is denoted y_ij_.

The EBP model is the nested error linear regression (NER) model (Battese, Harter & Fuller, 1988), which allows incorporation of both fixed and random effects to account for variability within and between domains. Additionally, it enables the use of unit-level (i.e., pixel-level) data, unlike area-level models (e.g., the Fay-Herriot model). The model is often used in finite-population models, as in this study (Sugasawa and Kubokawa, 2020). The NER model is expressed as:

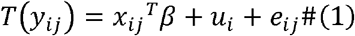

Where *T*(*y_ij_*) is the Box-Cox transformation of the dependent variable measured at the unit-level (pixel-level), *x*_*ij*_ is a p-vector of known covariates, with p the number of covariates, *β* is a p-vector of unknown fixed effects (the coefficient of regression), and *u_i_* and *e_ii_* are area-specific random effects and unit-level error terms, respectively. The underlying assumptions are that the fixed effect and the error term are mutually independent and distributed as *u_i_* ∼ 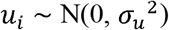 and 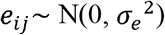. It is worth noting that, since *i_i_* is a domain-specific random effect and thus common to all the observations in the same domain, its variance 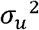 is often referred to as the “between” component of variance, in contrast to 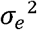 the “within” component.

Since the model assumes normality and homoscedasticity of the random effects and error terms, a transformation can be applied to relax these assumptions. The Box-Cox transformation is among the most widely used in practice, as the logarithmic transformation is a special case. The distribution of GSV volume within a sampling field dataset is already known to be right-skewed, both at the local and national level (Chirici et al., 2021; Vangi et al., 2022), so we transform the data using the Box-Cox transformation defined by:

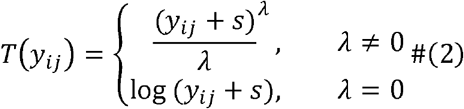

Where λ is an unknown transformation parameter; since the Box-Cox transformation needs positive values, s is a shift parameter that guarantees the non-negativity assumption. Both λ and s are estimated via the restricted maximum likelihood (REML) estimator (Pinheiro and Bates, 2002). Normality was assessed after transformation by inspecting Q-Q plots and using the Shapiro-Wilk (SW) and Kolmogorov-Smirnov (KS) tests (depending on the sample size). At the same time, homoscedasticity was evaluated by examining the skewness and excess kurtosis of residuals. Skewness and excess kurtosis values < |2| are considered acceptable for homoscedasticity (George & Mallery, 2010; Gravetter & Wallnau, 2014).

In model (2), the coefficients *β* and the variance components *u*_*i*_ + *e*_*ij*_ are estimated via the REML estimator. The EBP of small area parameters is computed as:

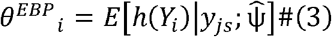

where Y_i_ = (y_1_, …,y_n_) are the unit values (both sampled and non-sampled), y_js_ are observed sample values in the j^th^ domain, 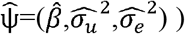 are model parameter estimates, h(·) is the functional form of the target parameter (linear or non-linear).

Monte Carlo approximation can be used to generate synthetic populations for the target variable, both for in-sample and out-of-sample domains, given x_ij_ and 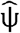, as explained in Kreutzmann et al. (2019). For more detailed information about the algorithm, refer to Kreutzmann et al. (2019), Molina and Rao (2010), and Molina and Marhuenda (2015). The small-area parameter for the i^th^ domain is estimated by averaging the Monte Carlo replicates’ parameter.

The MSE is estimated via a semi-parametric wild bootstrap procedure, described in detail by Kreutzmann et al. (2019, Appendix A) and Rojas-Perilla et al. (2019). Wild bootstrap not only takes into account the uncertainty derived from the transformation, but also protects against “‘mild’ departures from normality after using transformations” (Rojas-Perilla et al., 2019, p 123).

To express the estimator’s uncertainty on a meaningful scale, the square root of the MSE was computed, yielding the RMSE. The RMSE was also reported relative to the total and mean GSV for each stand, expressed as a percentage (RMSE%).

#### 2.5.2. The Mixed-effect Random Forests estimator

In Equation (1), the term 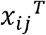 *β* models the conditional indicator (i.e., the mean) of y_*ij*_ given x_ij,_ but can be substituted with the general form *f (x*_*ij*_). While *f* is a linear mixed model in EBP, in MERF, *f* is a random forest model. The original procedure for estimating 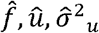 and 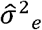 was developed by Hajjem et al. (2014) for random forests and called the Expectation–Maximization (EM) algorithm, which alternates between estimating the non-linear fixed effects and the random effects. The MERF algorithm, adjusted by Krennmair and Schmid (2022), fits Eq. 4 by iteratively estimating: 1) the forest function, assuming the correctness of random effects; 2) the random effect part, assuming the correctness of the out-of-bag (OOB) prediction of the random forest. Given for any small area i:

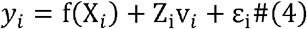

Where f (X_i_) represent any parametric or non-parametric function that expresses the conditional mean of y_i_ given X_i_ (random forest in the case of MERF), and Z_i_v_i_ is the linear part that captures the random effects. The iterative algorithm works as follows:

1. Set *b* = 0 and 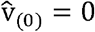
2. Set **b** = *b* + 1. Update 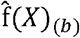 and 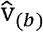 as follow:
  a. 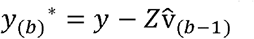
  b. Estimate 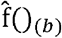 with random forest using *y* _(*b*)_* as the dependent variable and X as covariates
  c. Get the OOB predication from 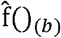
  d. Fit a linear mixed model with no intercept for 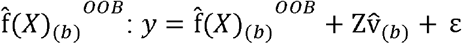
  e. Extract the variance component 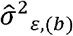 and the variance-covariance matrix 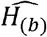 of *ε* and estimate the random effects by: 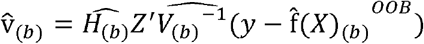
3. Repeat step 2) until convergence is reached. In this study, the convergence criterion is a marginal change in the generalized log-likelihood (GLL) of less than 1 ^−4,^ as already used by Krennmair and Schmid (2022) and Anand et al. (2024)

The estimation of the variance component 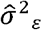 was naive and thus could not be considered a valid estimator for the variance of the unit-level error *ϵ*. Consequently, this estimator was corrected for bias to obtain the residual variance 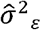 from the random forest model through the bootstrap approach proposed by Mendez and Lohr (2001).

A non-parametric random-effect block (REB) bootstrap is used to estimate the area-level MSE, following the formulation proposed by Chambers and Chandra (2013). The REB bootstrap ensures that the variability of residuals within each area is captured. The iterative algorithm for fitting eq (4) and the bootstrap MSE estimator are described in detail in Krennmair and Schmid (2022).

MERF was shown to be robust to model misspecification under EBP, particularly in the presence of non-Gaussian error structure and complex interactions among covariates.

Note that, since RF does not assume normality and normally distributed residuals, we fit Equation (4) without applying the Box-Cox transformation used in the EBP approach.

We used the SAEforest R package to fit the MERF algorithm, and we set the number of bootstrap iterations for MSE estimation to 200. We keep the default values for the RF hyperparameters, that is, ntree=500 and mtry=3.

#### 2.5.3. Validation

Both the NER and the RF model were evaluated using a buffer leave-one-out cross-validation (BLOOCV) scheme with the R package blockCV (Valavi et al., 2019), which creates evenly spatially distributed training and test sets (Figure S2), considering a buffer of a specified distance (ideally, the range of spatial autocorrelation) around each observation. Each fold is generated by excluding observations within the specified distance of each testing point, ensuring robust validation in the presence of possible spatial autocorrelation. The buffer distance (the range over which observations are independent) was determined by constructing the empirical variogram, a fundamental geostatistical tool for measuring spatial autocorrelation. The empirical variogram models the structure of spatial autocorrelation by measuring variability between all possible pairs of points (O’Sullivan and Unwin, 2010).

Models were evaluated using the mean R^2^, RMSE, and RMSE% across the BLOOCV iterations:

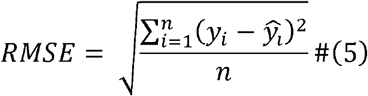

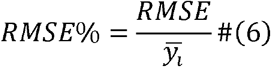

Where 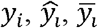 are the i^th^ observation of the dependent variable, the i^th^ prediction, and the mean of the observations, respectively; n represents the number of observations.

## 3. Results

### 3.1.1. model performance

The final NER model yielded a marginal R^2^ of 0.52 and a conditional R^2^ of 0.78 (including the random effect) on the training set. After back-transforming the data to the original range, the BLOOCV yielded an R^2^ of 0.37 and an RMSE of 202 m^3^ ha^-1^ (RMSE% = 28.9). The optimal λ and shift parameters for the Box-Cox transformation, estimated via the REML approach, resulted in λ = 0.45 and shift = 0, respectively. Normality of residuals, both for fixed and random effects, was assessed using the SW and KS tests, and by inspecting the QQ-plot (Figure S1). All tests indicated normality (p < 0.05). Homoscedasticity was evaluated by calculating the skewness and excess kurtosis of fixed and random residuals. Since skewness was 0.01 and 0.2 and excess kurtosis was 2.65 and 2.4 for fixed and random residuals, respectively, homoscedasticity was confirmed.

The MERF model achieved an R^2^ of 0.7 on the OOB observations. On the BLOOCV, the model achieved an R^2^ of 0.67 and an RMSE of 151 m^3^ ha^-1^, corresponding to 21.7% of the mean observed GSV. Figure 2 reports the results of the BLOOCV for both models, comparing predicted versus observed values.

**Figure 2.**
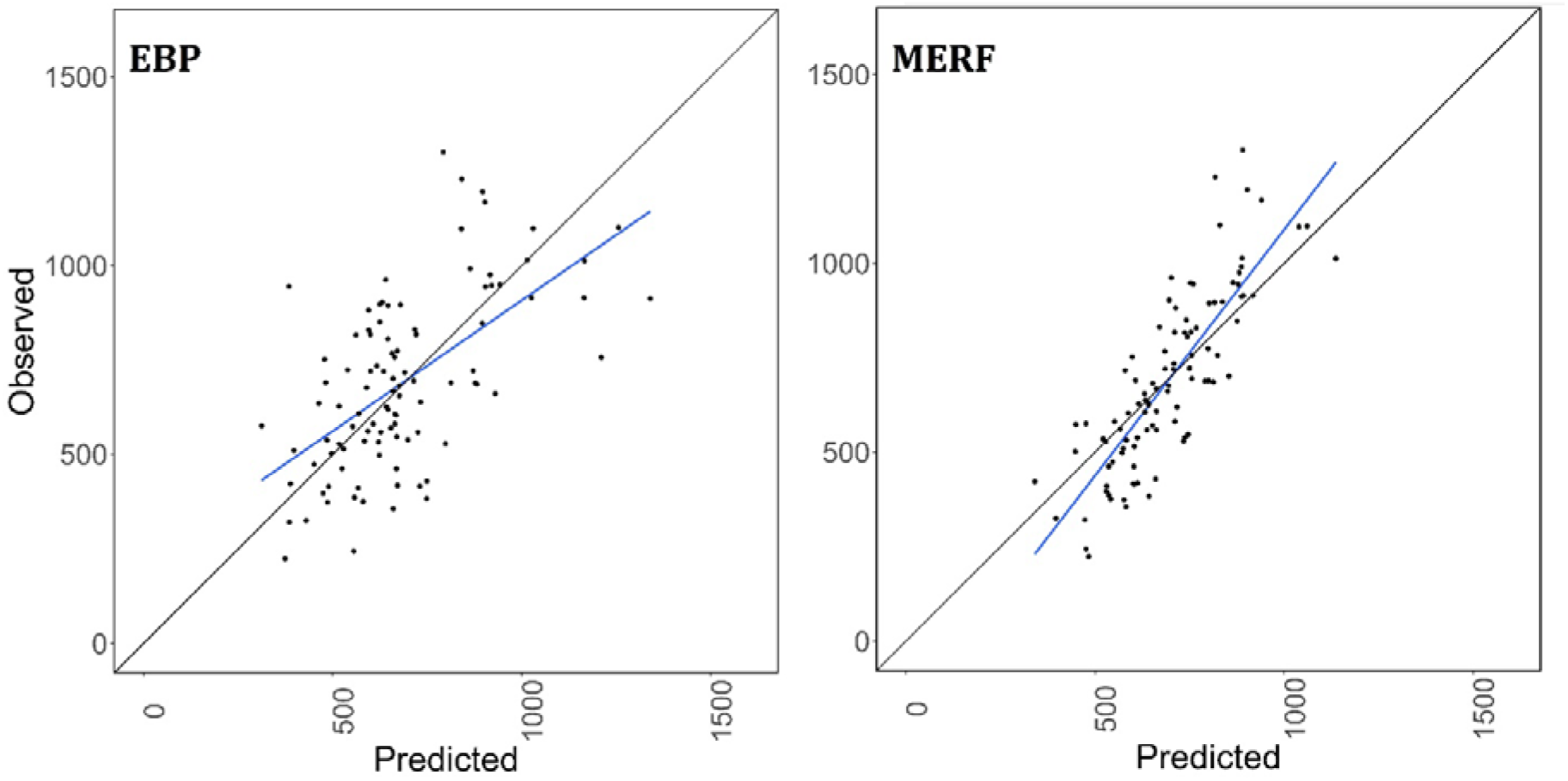
Scatterplots of predicted versus observed values for EBP (left panel) and MERF (right panel) resulting from the BLOOCV.

#### 3.2. Small area estimation

EBP estimation of mean GSV at the stand level ranged between 297 and 1821 m^3^ ha^-1^ with an average value of 767 m^3^ ha^-1^, while the mean GSV for MERF was between 350 and 1261 m^3^ ha^-1^ with an average of 761 m^3^ ha^-1^. The total GSV ranged from 181 and 219 m^3^ (minimum) to 8695 and 9995 m^3^ (maximum) for EBP and MERF, respectively. The average total GSV was 1768 and 1783 for EBP and MERF, respectively.

The RMSE for the mean GSV varies from 54 to 1768 with a mean of 198 m^3^ ha^-1^ (EBP) and from 51 to 197 with a mean of 155 m^3^ ha^-1^ (MERF).

The observed mean of the sampled domains fell within the corresponding SAE interval (1.96*RMSE) for 66 of 79 sampled domains for EBP and 77 of 79 for MERF (Fig. 3).

**Figure 3.**
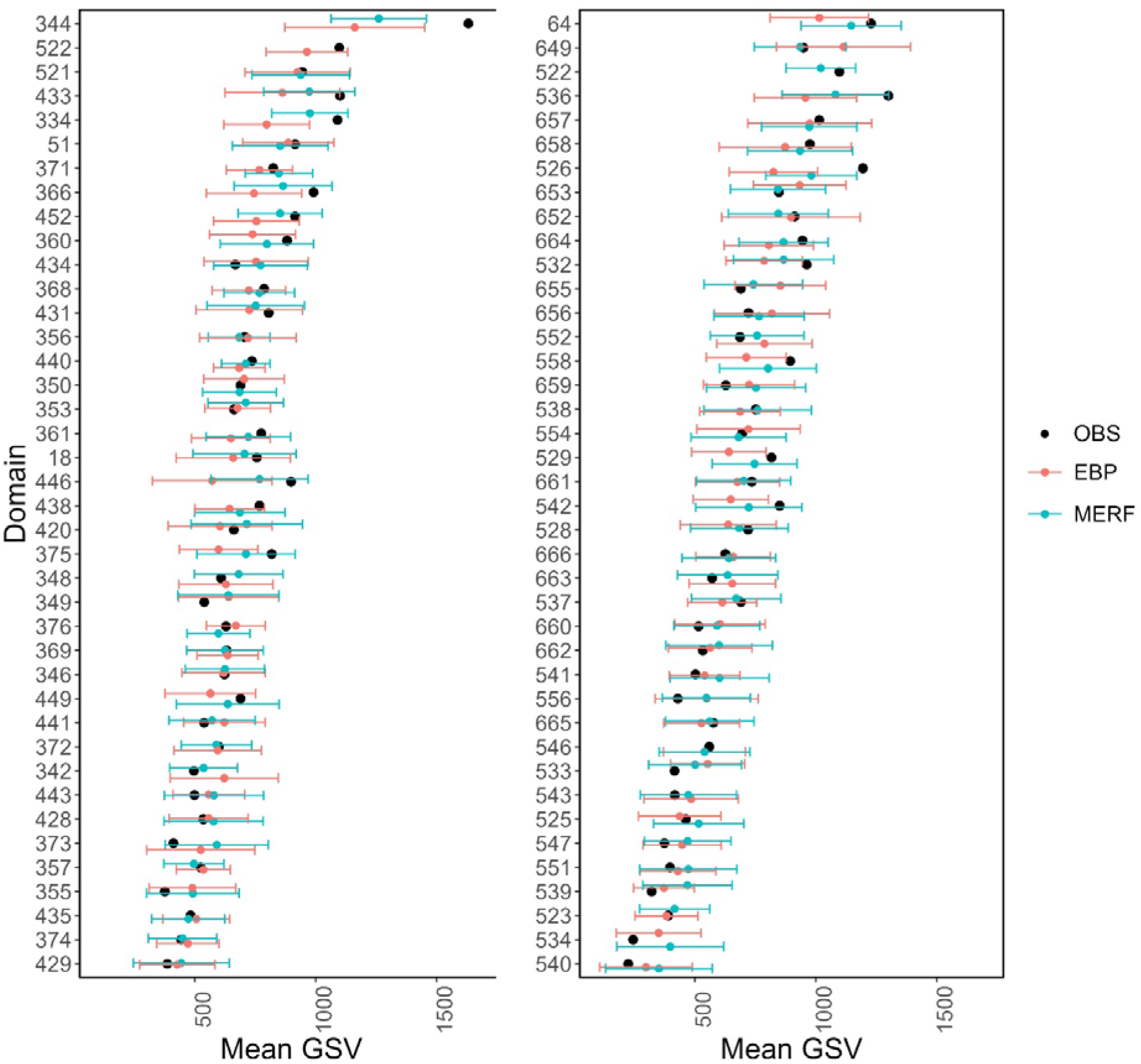
Estimate of mean GSV and its RMSE at the domain level obtained with EBP and MERF. The black dot represents the mean GSV from the observed sampling point.

Table 2 summarizes the results for the mean and total GSV, along with their respective RMSEs, for both EBP and MERF.

**Table 2.** Results for mean and total GSV and their respective RMSE for EBP and MERF models.

|  |  | Mean GSV |  | Total GSV |  |
| --- | --- | --- | --- | --- | --- |
|  |  | SAE | RMSE | SAE | RMSE |
| EBP | min | 297 | 54 | 1814 | 43 |
|  | mean | 767 | 198 | 1768 | 470 |
|  | max | 1821 | 1768 | 8695 | 4119 |
|  | sd | 225 | 126 | 1237 | 463 |
| **MERF** | min | 351 | 51 | 219 | 48 |
|  | mean | 761 | 155 | 1783 | 366 |
|  | max | 1261 | 197 | 9995 | 2736 |
|  | sd | 110 | 25 | 1257 | 275 |

Figures 4 and 5 show the mean GSV value at the forest stand level (left panel) and its standard error (right panel) for EBP and MERF, respectively, while Figure 6 shows the distribution of mean and total GSV, along with their RMSE and RMSE%.

**Figure 4.**
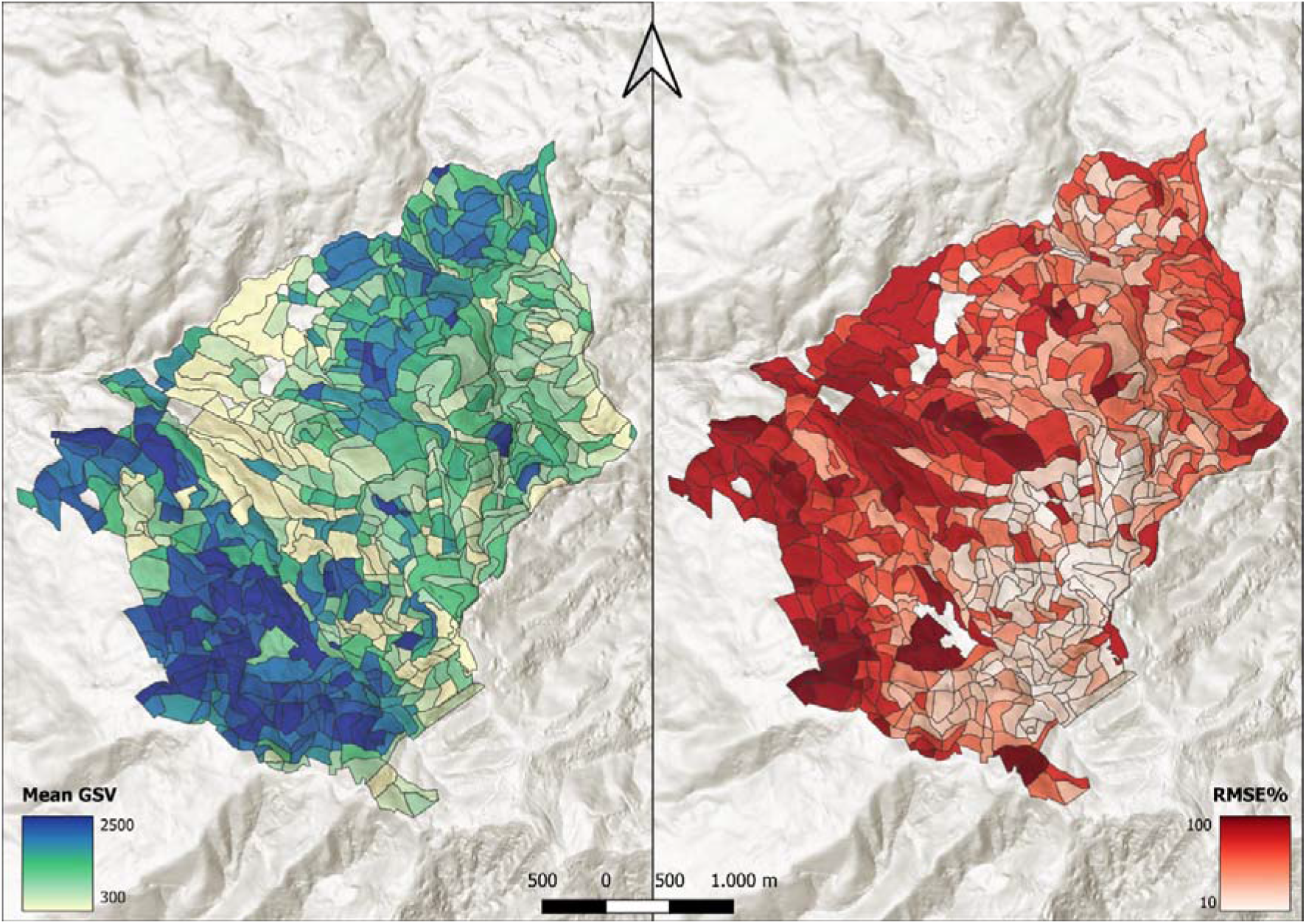
Map of mean GSV (left panel) and its RMSE% (right panel) using the EBP estimator at the forest stand level.

**Figure 5.**
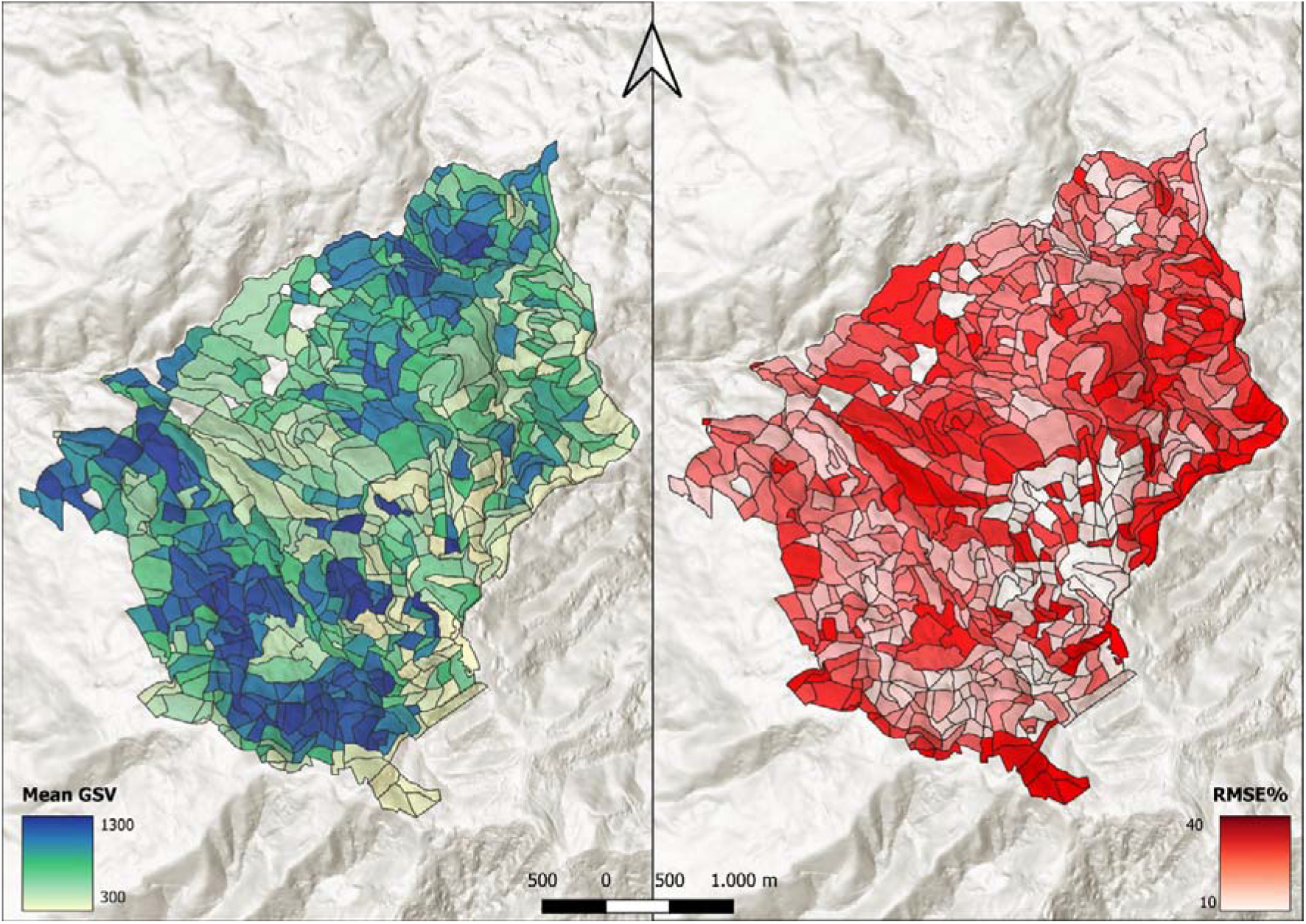
Map of mean GSV (left panel) and its RMSE% (right panel) using the EBP estimator at the forest stand level.

**Figure 6.**
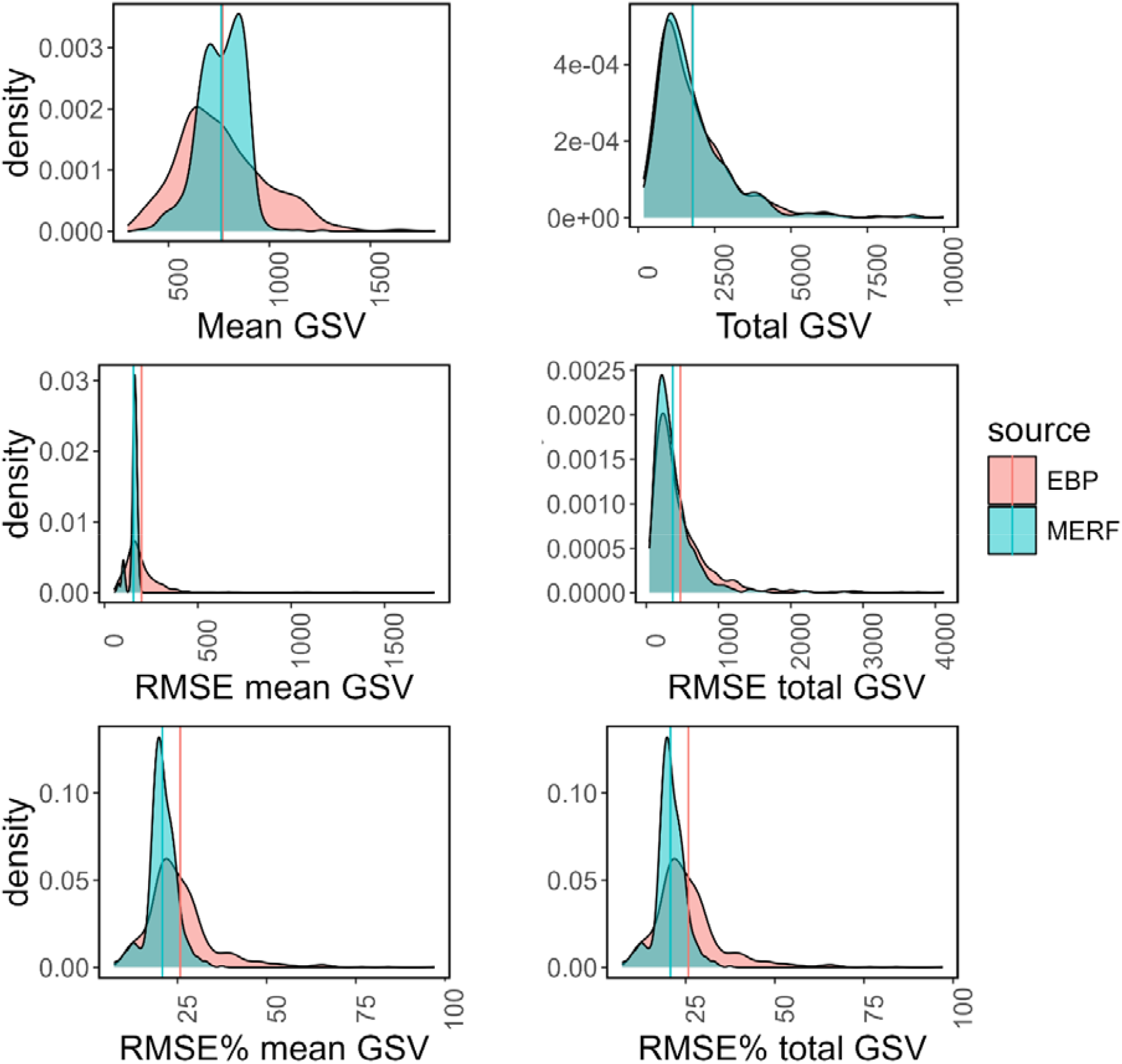
Distribution of mean and total SAE and their respective RMSE and RMSE% for EBP (red) and MERF (blue). The vertical lines represent the mean values.

The RMSE of the mean GSV decreases with increasing plot count within the domain, but not linearly (Fig. S3), for both methods. In contrast, the RMSE of the total GSV varies with the domain dimension rather than with the number of sampling points. Overall, the RMSE of MERF for mean and total shows a narrower distribution around the mean than EBP.

A positive relationship between GSV and its RMSE is observed in mean and total estimates for both methods, but with significant differences between the two approaches (Fig. 7). EBP shows a higher standard deviation and a steeper regression line. Furthermore, several outliers are observed for high GSV values (over ∼1200 m^3^ ha^-1^), with RMSE% approachin 100%. In contrast, MERF shows a more compact point distribution and a much shallower slope. The RMSE remains relatively stable across the entire GSV range, with lower variability and fewer extreme outliers than observed for EBP.

**Figure 7.**
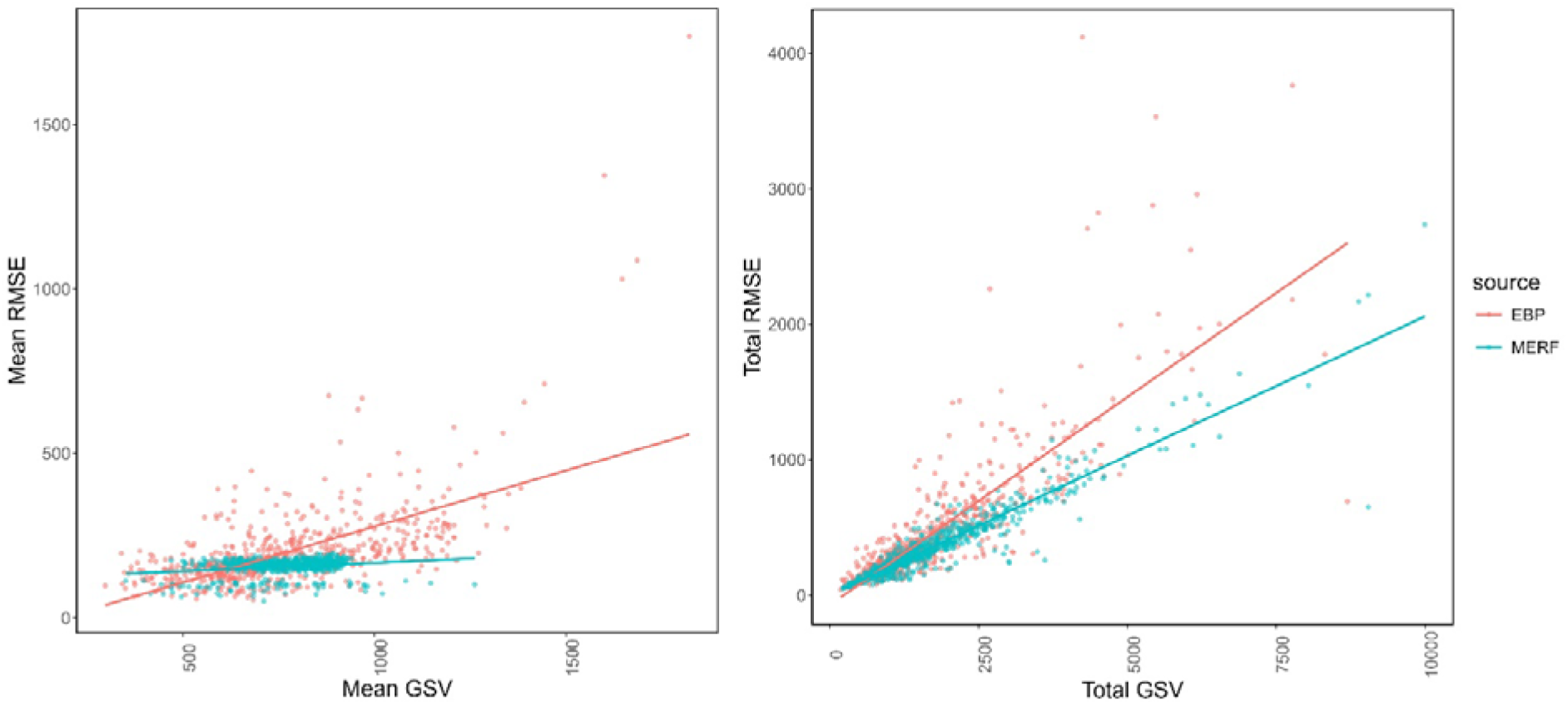
Scatterplot of mean (left panel) and total (right panel) GSV against their respective RMSE. Each dot represents a single domain (forest stand)

## 4. Discussion

Direct estimates of mean and total GSV at the forest stand level are limited to sampled domains and, therefore, unavailable for all areas of interest. The use of model-based SAE methods that incorporate unit-level auxiliary information enables extending predictions to non-sampled domains while improving overall estimate precision compared to direct estimators or even to area-level mixed models, such as the Fay–Herriot model (Fay & Herriot, 1979). Numerous forestry studies have demonstrated the advantage of using indirect or composite estimators over direct, design-based estimators (e.g., Breidenbach et al., 2012, 2018; Ver Plank et al., 2018; Mauro et al., 2017, 2019; Frank et al., 2020; Georgakis et al., 2025; see also the review by Dettman et al., 2021). The resulting maps reveal gradients and clusters of forest volume that would not be observable using sampled data alone, highlighting the practical value of SAE methods in forest inventory and management. Given the skewed distribution of GSV, model misspecification was a potential concern when using standard linear mixed models. For this reason, both the EBP approach with dependent variable transformation and the MERF model were considered appropriate alternatives for handling non-normal data, as confirmed by the results of the assumption tests.

The comparison between EBP and MERF highlights important differences in predictive performance, robustness, and suitability for SAE of GSV. While both approaches enable the extension of estimates to non-sampled forest stands and provide comparable estimates of total GSV, their behavior differs substantially in terms of accuracy, variability, and stability across domains.

The NER model showed a moderate explanatory power on the training data, with a marginal R^2^ of 0.52 and a conditional R^2^ of 0.78, confirming the importance of random effects in capturing hierarchical variability. However, this improvement did not fully translate into predictive performance, as indicated by the lower BLOOCV R^2^ (0.37) and relatively high RMSE (202 m^3^ had^1^; 28.9%). Although the Box–Cox transformation ensured compliance with model assumptions, these statistical properties did not yield strong generalization performance. This suggests that, despite being well-specified, the NER model is unable to capture complex, non-linear relationships under heterogeneous forest conditions (White et al., 2021; May et al., 2023). Moreover, multicollinearity in the training dataset may inflate the variance of the regression coefficients, thereby affecting unit-level predictions and the variance component of the MSE estimator (Rao and Molina, 2015).

In contrast, the MERF model demonstrated consistently greater predictive performance, achieving R^2^ values of 0.70 (OOB) and 0.67 under BLOOCV, along with lower RMSE (151 m^3^ ha^-1^; 21.7%). These results indicate stronger generalization capability, likely due to MERF’s ability to model non-linear relationships and interactions among predictors while accounting for hierarchical structure through random effects. Furthermore, MERF is more robust to multicollinearity due to its underlying RF model, which prevents variance inflation and ultimately yields more stable MSE estimates. Anand et al. (2024) estimated per capita expenditure in Indonesia, demonstrating that MERF effectively handles multicollinearity, especially in very small domains. This advantage becomes particularly relevant in forest ecosystems, where structural complexity and environmental variability often violate the assumptions of parametric models, and multicollinearity among remote sensing variables is common. Partial dependence plot in supplementary materials (Fig. S4) reveals the complex, non-monotonic, non-linear relations between GSV and almost all covariates, except for the CHM.

At the small-area level, both methods produced slightly overestimated overall mean GSV estimates (767 m^3^ ha^-1^ for EBP and 761 m^3^ ha^-1^ for MERF against an observed mean of 707 m^3^ ha^-1^), but with different distributions of the domain’s means. In particular, the RF predictions tend to cluster around the observed mean, a phenomenon known as shrinkage towards the mean. This behavior is consistent with the properties of random forest, which introduce implicit regularization through variance reduction via ensemble averaging (Breiman, 2001; Hastie et al., 2009). Also, Krennmair and Schmid (2020) observed the same effect when estimating total household per capita income in Mexico, using the MERF model. The same phenomenon had already been observed in biomass and carbon mapping, confirming that the shrinkage effect is not an emergent feature of this study. For example, Zhang et al. (2024) found that regularized RF and quantile RF significantly reduce overestimation and underestimation compared with vanilla RF. Mascaro et al. (2014) found that adding spatial context to RF improved tropical forest carbon mapping. The reduction in prediction dispersion lowers the overall RMSE, but at the cost of reduced ability to accurately represent extreme values. The shrinkage effect implies that the model underestimates high GSV values and overestimates low ones, as suggested by the weak correlation between RMSE and GSV. This is not a MERF-specific feature, despite being amplified by the RF implicit regularization; in Kubokawa’s (2009) and Kubokawa’s and Sugasawa’s (2020) reviews on SAE, the authors point out that among the desirable properties of the EBLUP are the shrinkage function and the pooling effects, which arise from the setup of random effects and common parameters in LMM, indicating the intrinsic shrinkage behavior of the SAE framework.

The same effect is not observed in the total GSV estimate, which instead shows strong agreement with the EBP method. This apparent discrepancy can be explained by the scale-dependent behavior of the bias–variance trade-off. At the unit level, MERF reduces variance through ensemble averaging, resulting in smoother predictions and a tendency to shrink extreme values toward the mean. When unit-level predictions are summed to obtain area-level totals, individual over- and under-estimations tend to compensate, and the variance reduction achieved by shrinkage dominates over the bias (Molina and Rao, 2015), leading to more consistent total estimates. Future studies should focus on mitigating the shrinkage effect of RF by exploring, e.g., post-hoc bias correction, hyperparameterization, and estimating the conditional distribution rather than the mean, e.g., with quantile random forest regression.

More pronounced differences emerge when considering uncertainty measures. The RMSE associated with MERF predictions is consistently lower and exhibits a narrower distribution (mean RMSE = 155 m^3^ ha^-1^) compared to EBP (198 m^3^ ha^-1^), indicating greater precision and stability. Additionally, the coverage of observed values within the SAE confidence intervals is higher for MERF (77 out of 79 domains) than for EBP (66 domains), suggesting improved reliability in uncertainty quantification. However, it is worth noting that the size of small areas of interest plays a fundamental role in determining the precision of estimates within the SAE framework. Although SAE techniques are designed to produce reliable estimates at increasingly fine spatial resolutions, the number of sample plots available within each area (*n*_*d*_) remains a critical factor. Specifically, the sampling variance of direct estimators in inversely proportional to *n*_*d*_, implying that small sample sizes lead to high variability and unstable estimates. SAE methods mitigate this limitation by leveraging auxiliary information; however, this introduces a trade-off between variance reduction and potential model-induced bias. (Molina and Rao, 2015; Georgakis et al., 2025). In particular, in unit-level methods such as NER models under EBP and RF under MERF, the MSE comprises sampling variance, model variance, and parameter uncertainty. When *n*_*d*_ is small, estimates rely more heavily on model predictions, leading to stronger shrinkage and MSE dominated by model assumptions. As *n_d_* increases, the influence of direct observations grows, reducing shrinkage and lowering MSE. Our study was characterized by low coverage of the sampling domain (79 out of 658) and low sampling intensity within domains, with a mean of 1 sampling point per domain and a maximum of 4, suggesting a strong dependence of the estimate MSE on the model prediction. In turn, this dependency confirms the goodness-of-fit of both models, given the relatively low RMSE obtained over the domains. Rahalf et al. (2014) obtained similar performance (RMSE%=18.2) in estimating timber volume within small forest stands (1-3 ha) using the NER model, but with a higher sampling intensity of 5-7 plots per domain.

The behavior of RMSE in relation to domain characteristics provides further insight. For both methods, the RMSE of mean GSV decreases with increasing sample size, although not linearly, reflecting diminishing returns from increased sampling intensity. In contrast, the RMSE of total GSV is more strongly influenced by domain size than by the number of sampling points, emphasizing the role of spatial extent in error propagation. A positive relationship between total GSV and RMSE is observed for both approaches; however, this relationship is not observed for the mean estimate in the MERF approach, while it still exists for EBP, which shows higher variability and several extreme outliers, particularly for high GSV values (above ∼1200 m^3^ ha□^1^), where RMSE% approaches 100%. This difference in behavior between EBP and MERF can be attributed to their underlying model structures: In the NER model, the back transformation inherently links the variance of predictions to the magnitude of the estimated variable, leading to a proportional increase in RMSE with both the mean and total estimates (Molina and Rao, 2010). In contrast, MERF does not impose a parametric relationship between the mean and variance, as prediction uncertainty is driven by the residual structure of the random forest and random effects. Consequently, the RMSE of the mean does not necessarily increase with the magnitude of the estimate, although the RMSE of totals still scales with the population size (domain’s area) due to aggregation. It should be noted, however, that EBP has a mathematically proven estimator of MSE, firmly grounded in statistical theory. At the same time, the non-parametric bootstrap scheme of MERF remains an active area of statistical research regarding its theoretical consistency in small domains (Krennmair and Schmit., 2022).

From a modeling perspective, a key distinction lies in how predictors are treated. The EBP approach requires explicit model specification and variable selection, whereas MERF benefits from the implicit variable selection of random forests, enabling it to identify relevant predictors, capture complex interactions, and reduce preprocessing steps. While this improves predictive performance, it also introduces a trade-off in interpretability (Krennmair and Schmid, 2020). Nonetheless, interpretability is often less important than estimate precision in the SAE framework. Interpretability can be partially recovered through diagnostic tools such as variable importance and partial dependence analysis, as well as explainable AI methods such as SHapley Additive exPlanations (SHAP) (Stumbelj et al., 2014). However, it must be noted that the computational time required to fit the MERF model was more than six times that of the EBP, especially due to the complex non-parametric bootstrap scheme for RMSE estimation and the bootstrap-based adjustment of the residual variance in RF. For large-scale applications, computational intensity can become an issue, which could be partially mitigated by parallelizing the bootstrap scheme.

Overall, the results confirm that model-based SAE methods substantially enhance the estimation of forest attributes at fine spatial scales. While the EBP framework remains theoretically robust and interpretable, its predictive performance is limited in complex and highly variable forest environments. The MERF model, by combining the strengths of machine learning and mixed-effects modeling, provides a more accurate, stable, and reliable alternative for GSV estimation. These findings support the adoption of machine learning-based approaches in forest inventory, particularly for handling non-linear relationships, heterogeneous conditions, and low-intensity sampling coverage.

## 5. Limitation and future direction

Forest variables are inherently spatial. While SAE models include area-level random effects, traditional RF and NER models do not explicitly model continuous spatial autocorrelation. To mitigate spatial autocorrelation, we included the coordinates and altitude of the sampling points as predictors and evaluated the model using a spatial block cross-validation approach. However, future studies should focus on integrating explicit spatial models into SAE, for example, comparing the spatial empirical best linear unbiased predictor (SEBLUP, Petrucci and Salvati, 2004a; 2004b) or the spatial empirical Bayes predictor (SEBP, Handayani et al., 2018) and including Geographically Weighted RF or Spatial RF into MERF. Emick et al. (2023) proposed a Geostatistical model-based estimator to account for spatial autocorrelation and to correct for bias, based on summaries of Monte Carlo Markov Chain samples from Bayesian posterior distributions. Another possibility to relax the NER model’s assumptions, particularly in the presence of multicollinearity or outliers, is to consider robust estimation techniques, such as the estimators developed by Chambers et al. (2014) and further extended by Schmid et al. (2016) to account for spatial correlation. On the other hand, Ananda et al. (2023) have already used Principal Component Analysis and rotation forests in MERF to mitigate multicollinearity.

Regarding the indicators, while the mean is the gold standard in almost all SAE research, totals and other indicators are often understudied, especially indirect indicators (indicators that are not in a linear relationship with the unit-level predictions), such as the Gini index or quantiles, which may be interesting parameters in biodiversity conservation studies.

Finally, an inherent limitation, and, at the same time, a notable feature of the study, was the large proportion of unsampled domains and the small sampling size within domains (n_*d*_ ≤ 4), which limited the accuracy of the estimation in terms of RMSE, but highlighted the necessity of such approaches to deal with real-world datasets, often with similar charateristics in the field of local forest management.

## 6. Conclusion

Small-area estimation has become an increasingly important research focus in forest inventory, primarily because it enables model-based inference in small domains where limited sample data prevent reliable estimates based solely on design-based methods, and secondly because of its inherent ability to provide an assessment of the estimate via standard error or MSE estimation. While the EBP approach remains theoretically sound and interpretable, its performance is constrained by complex, non-linear relationships and multicollinearity in the training dataset. In contrast, the MERF model offers advantages in model specification, affecting its predictive accuracy, robustness, and stability, providing more reliable uncertainty estimates, albeit at the cost of increased computational demand and reduced interpretability. Both methods exhibit inherent shrinkage effects that influence domain-level estimates but tend to balance out at aggregate levels, resulting in consistent total GSV estimates. Ultimately, the findings highlight a trade-off between theoretical rigor and predictive performance, supporting the integration of machine-learning-based approaches into forest inventory practices and local management plans, especially in contexts characterized by low sampling intensity and high environmental heterogeneity, thereby contributing to the ongoing development of more robust SAE methods.

## Supporting information

appendix of the main manuscript

## Availability of data and material

Sentinel-2 data are freely available at the original source. All other data and codes will be shared upon request to the author.

## Acknowledgments

E.V. acknowledges NextGenCarbon H2020 project funded by the European Commission, number 101184989 call HORIZON-CL5-2024-D1-01-07 and the Space It Up! project funded by the Italian Space Agency, ASI, and the Ministry of University and Research, MUR, under contract n. 2024-5-E.0 - CUP n. I53D24000060005.

## Appendix

In this section, the NER model’s performance and other meaningful statistical metrics are reported, as well as the spatial scheme of the BLOOCV, the relationship between RMSE and sampling intensity and the partial dependence plot for the RF model.

**Figure S1.**
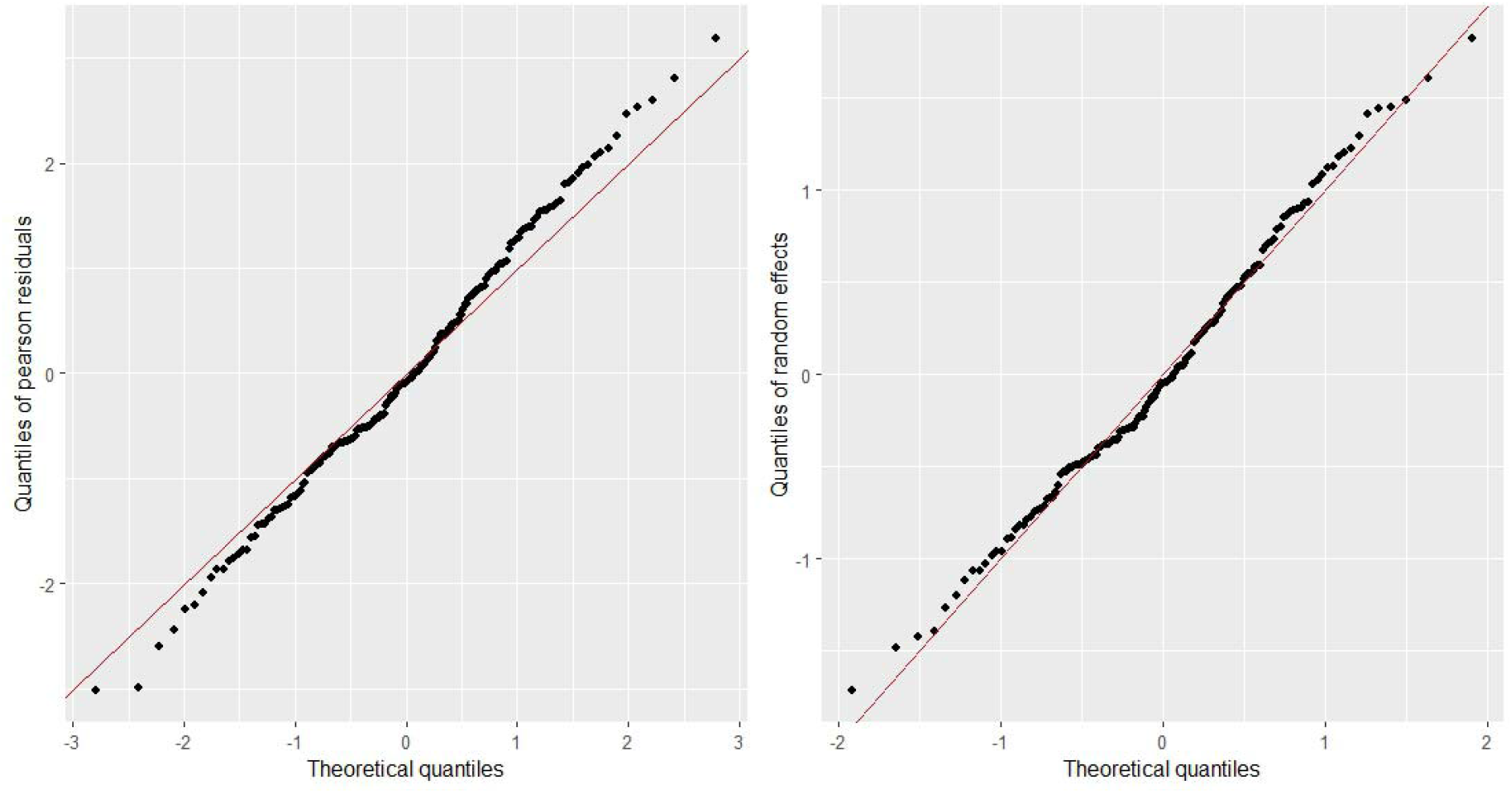
QQ-plot of fixed (left panel) and random (right panel) residuals.

**Figure S2.**
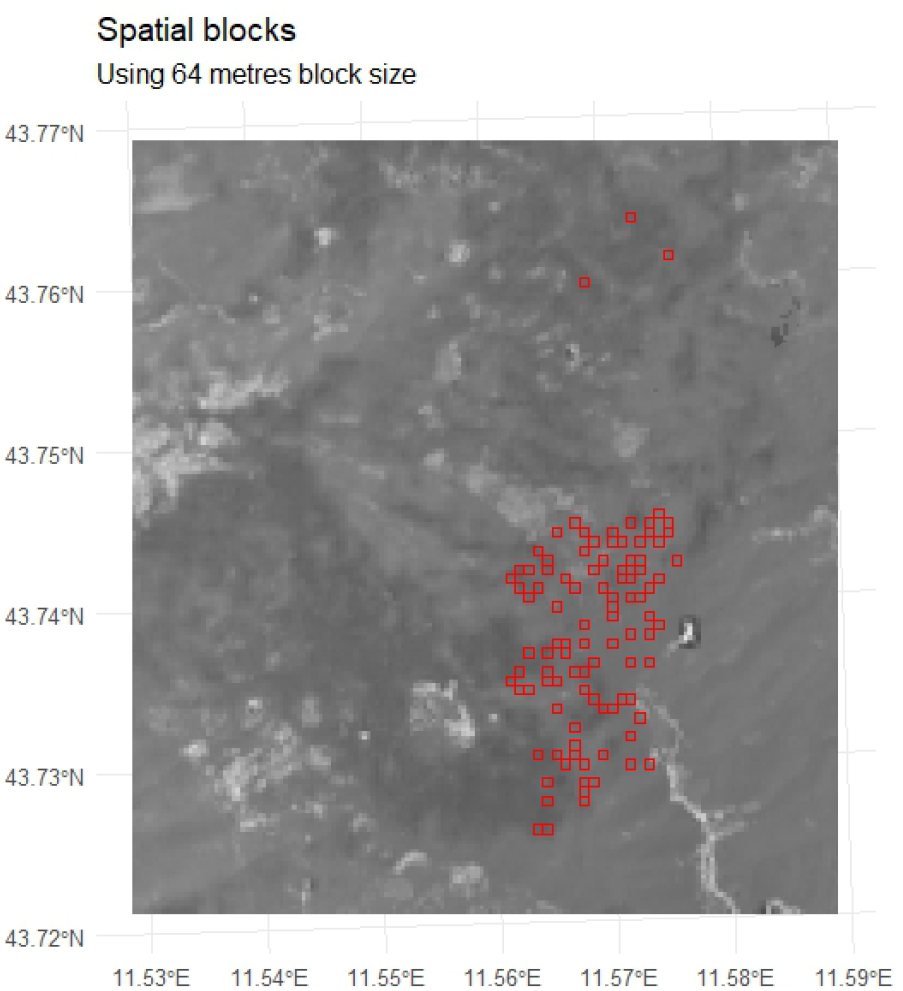
Spatial distribution of the 10 blocks generated for the cross-validation.

**Figure S3.**
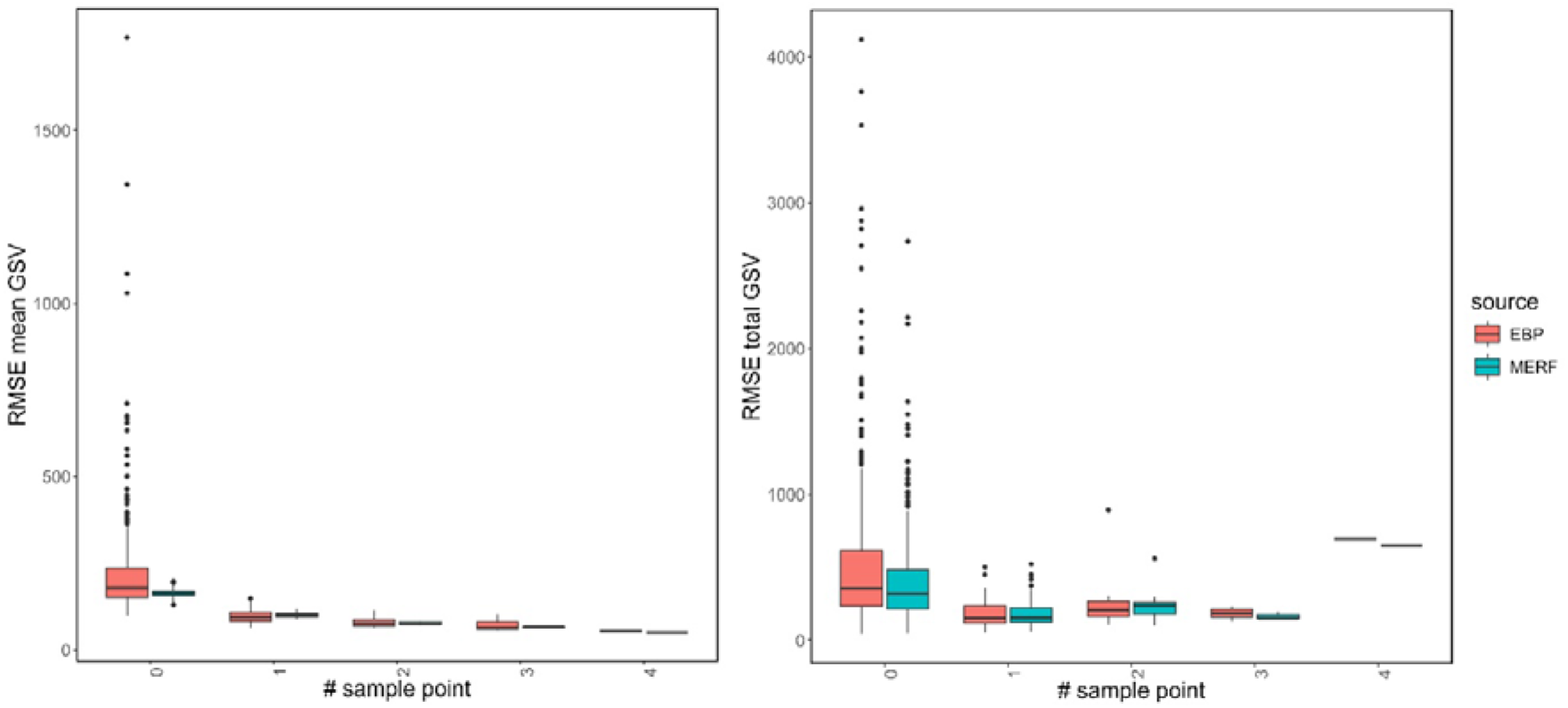
Distribution of RMSE for mean (left panel) and total (right panel) GSV at varying numbers of sampling plots within domains.

**Figure S4.**
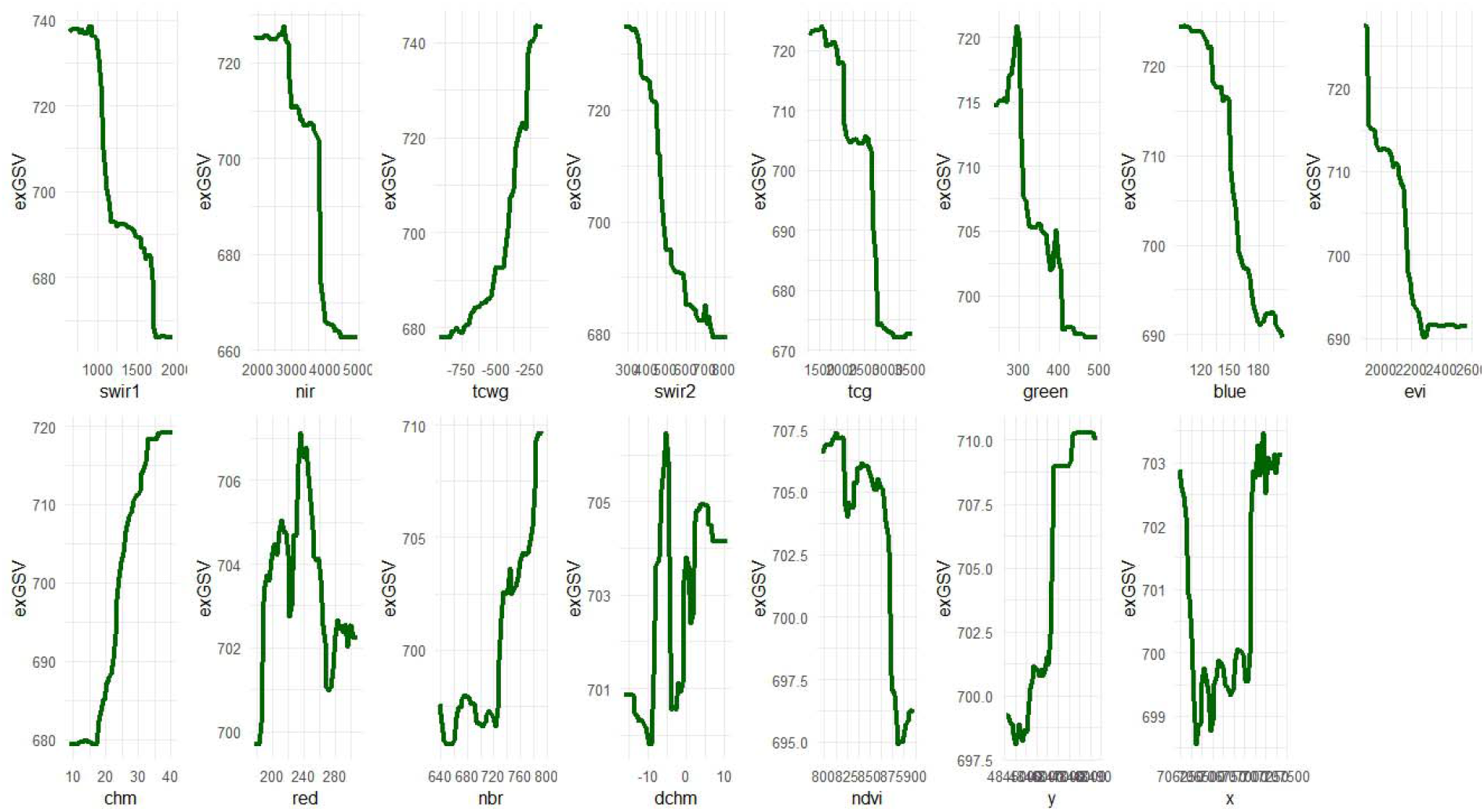
Partial dependence plot for variables ranked by their permutation importance

## References

Ananda, R., Notodiputro, K. A., & Aidi, M. N. (2024). Modified Mixed Effects Random Forest in Small Area Estimation Using PCA and Rotation Forest with Correlated Auxiliary Variables. Scientific Journal of Informatics, 11(3), 705–720.

Bajocco, S., Ferrara, C., Alivernini, A., Bascietto, M., & Ricotta, C. (2019). Remotely-sensed phenology of Italian forests: Going beyond the species. International Journal of Applied Earth Observation and Geoinformation, 74, 314–321.

Battese GE, Harter RM, Fuller WA (1988). “An Error-Components Model for Prediction of County Crop Areas Suing Survey and Satellite Data.” Journal of the American Statistical Association, 83(401), 28–36. doi:10.1080/01621459.1988.10478561.

Breiman L (2001). “Random Forests.” Machine Learning, 45(1), 5–32.

Breidenbach, J., & Astrup, R. (2012). Small area estimation of forest attributes in the Norwegian National Forest Inventory. European Journal of Forest Research, 131(4), 1255–1267.

Breidenbach, J., Magnussen, S., Rahlf, J., & Astrup, R. (2018). Unit-level and area-level small area estimation under heteroscedasticity using digital aerial photogrammetry data. Remote Sensing of Environment, 212, 199–211.

Chambers R, Chandra H (2013). “A Random Effect Block Bootstrap for Clustered Data.” Journal of Computational and Graphical Statistics, 22(2), 452–470.

Chambers, R., Chandra, H., Salvati, N., & Tzavidis, N. (2014). Outliner robust small area estimation. Journal of the Royal Statistical Society: Series B, 76, 47–69.

Chirici, G., Giannetti, F., McRoberts, R.E., Travaglini, D., Pecchi, M., Maselli, F., Chiesi, M., Corona, P., 2020. Wall-to-wall spatial prediction of growing stock volume based on Italian National Forest Inventory plots and remotely sensed data. Int. J. Appl. Earth Obs. Geoinf. 84, 101959 10.1016/J.JAG.2019.101959.

Ciancio, O. Riserva Naturale Statale Biogenetica di Vallombrosa. Piano di Gestione e Silvomuseo 2006–2025; Corpo Forestale dello Stato,Ufficio Territoriale per la Biodiversità di Vallombrosa, Reggello (FI): Florence, Italy, 2009; pp. 113–134. ISBN 978-88-87553-17-8.

D’Amico, G., Francini, S., Giannetti, F., Vangi, E., Travaglini, D., Chianucci, F., Mattioli, W., Grotti, M., Puletti, N., Corona, P., Chirici, G., 2021. A deep learning approach for automatic mapping of poplar plantations using S2 imagery. Giscience & Remote Sensing 58 (8), 1352–1368. 10.1080/15481603.2021.1988427.

D’Amico, G., Chirici, G., Corona, P., Romano, R., Di Domenico, G., Giannetti, F., & Mattioli, W. (2023). Differenze locali e prospettive globali per le foreste italiane: La definizione di bosco nel prossimo Sistema Informativo Forestale Nazionale. L’Italia forestale e montana, 78(1), 15–29.

Dettmann, G. T., Radtke, P. J., Coulston, J. W., Green, P. C., Wilson, B. T., & Moisen, G. G. (2022). Review and synthesis of estimation strategies to meet small area needs in Forest inventory. Frontiers in Forests and Global Change, 5, 813569.

Emick, E., Babcock, C., White, G. W., Hudak, A. T., Domke, G. M., & Finley, A. O. (2023). An approach to estimating forest biomass while quantifying estimate uncertainty and correcting bias in machine learning maps. Remote Sensing of Environment, 295, 113678.

Fay III, R. E., & Herriot, R. A. (1979). Estimates of income for small places: an application of James-Stein procedures to census data. Journal of the American Statistical Association, 74(366a), 269–277.

Foody, G.M., Boyd, D.S., Cutler, M.E.J., 2003. Predictive Relations of Tropical Forest Biomass from Landsat TM Data and Their Transferability between Regions 85. pp. 463–474. 10.1016/S0034-4257(03)00039-7.

Frank, B. M. (2020). Aerial laser scanning for forest inventories: estimation and uncertainty at multiple scales. PhD diss Oregon State University, 2020

Georgakis, A., Papageorgiou, V. E., & Stamatellos, G. (2025). A new approach to small area estimation: Improving forest management unit estimates with advanced preprocessing in a multivariate Fay–Herriot model. Forestry: An International Journal of Forest Research, 98(4), 605–622.

George, D., & Mallery, M. (2010). SPSS for Windows Step by Step: A Simple Guide and Reference, 17.0 update (10a ed.) Boston: Pearson

Giannetti, F.; Chirici, G.; Gobakken, T.; Naesset, E.; Travaglini, D.; Puliti, S. A new approach with DTM-independent metrics for forest growing stock prediction using UAV photogrammetric data. Remote Sens. Environ. 2018, 213, 195–205.

Goerndt, M. E., Monleon, V. J., & Temesgen, H. (2013). Small-area estimation of county-level forest attributes using ground data and remote sensed auxiliary information. Forest Science, 59(5), 536–548.

Gorelick, N., Hancher, M., Dixon, M., Ilyushchenko, S., Thau, D., and Moore, R. (2017). Google earth Engine: Planetary-scale geospatial analysis for everyone. Remote Sens. Environ. 202, 18–27. doi: 10.1016/j.rse.2017.06.031

Gravetter, F., & Wallnau, L. (2014). Essentials of statistics for the behavioral sciences (8th ed.). Belmont, CA: Wadsworth.

Hajjem A, Bellavance F, Larocque D (2014). “Mixed-Effects Random Forest for Clustered Data.” Journal of Statistical Computation and Simulation, 84(6), 1313–1328.

Handayani, D., Folmer, H., Kurnia, A. et al. The spatial empirical Bayes predictor of the small area mean for a lognormal variable of interest and spatially correlated random effects. Empir Econ 55, 147–167 (2018). 10.1007/s00181-018-1452-5

Hobza, T., & Morales, D. (2016). Empirical best prediction under unit-level logit mixed models. Journal of official statistics, 32(3), 661–692.

Jönsson, P., Cai, Z., Melaas, E., Friedl, M. A., & Eklundh, L. (2018). A method for robust estimation of vegetation seasonality from Landsat and Sentinel-2 time series data. Remote Sensing, 10(4), 635.

Kennedy, R.E., Yang, Z., Gorelick, N., Braaten, J., Cavalcante, L., Cohen, W.B., Healey, S., 2018. Implementation of the landtrendr algorithm on google earth engine. Remote Sens. 10, 1–10. 10.3390/rs10050691.

Krennmair, P. Tree-Based Machine Learning in Small Area Estimation, 10 (2022).

Krennmair, P., Würz, N. & Schmid, T. Analysing Opportunity Cost of Care Work using Mixed Effects Random Forests under Aggregated Census Data Apr. 22, 2022. 2204.10736[stat]. http://arxiv.org/abs/2204.10736 (2022). 37.

Krennmair, P. & Schmid, T. Flexible domain prediction using mixed effects random forests. Journal of the Royal Statistical Society: Series C (Applied Statistics). 2201.10933 [stat], rssc.12600. issn: 0035-9254, 1467-9876. http://arxiv.org/abs/2201.10933 (2022) (Oct. 2022)

Kreutzmann AK, Pannier S, Rojas-Perilla N, Schmid T, Templ M, Tzavidis N (2019). emdi: Estimating and Mapping Disaggregated Indicators. R package version 1.1.6, URL https://CRAN.R-project.org/package=emdi.

Kubokawa, T. (2009). A Review of Linear Mixed Models and Small Area Estimation. CIRJE F-Series.

Magnussen, S., & Breidenbach, J. (2017). Model-dependent forest stand-level inference with and without estimates of stand-effects. Forestry: An International Journal of Forest Research, 90(5), 675–685.

Mascaro, J., Asner, G. P., Knapp, D. E., Kennedy-Bowdoin, T., Martin, R. E., Anderson, C., … & Chadwick, K. D. (2014). A tale of two “forests”: Random Forest machine learning aids tropical forest carbon mapping. PloS one, 9(1), e85993.

Mauro, F., Monleon, V. J., Temesgen, H., & Ford, K. R. (2017). Analysis of area level and unit level models for small area estimation in forest inventories assisted with LiDAR auxiliary information. PloS one, 12(12), e0189401.

McRoberts, R., Tomppo, E., Schadauer, K., Vidal, C., Stahl, G., Chirici, G., Lanz, A., Cienciala, E., Winter, S., Smith, B., 2009. Harmonizing national forest inventories. J. For. 179–187.

Mendez, G., and Lohr, S., “Estimating residual variance in random forest regression,” Computational statistics & data analysis, vol. 55, no. 11, pp. 2937–2950, 2011, doi: 10.1016/j.csda.2011.04.022.

Molina I, Marhuenda Y (2015). “sae: An R Package for Small Area Estimation.” The R Journal, 7(1), 81–98. doi:10.32614/rj-2015-007.

Molina I, Martín N (2018). “Empirical best prediction under a nested error model with log transformation.” The Annals of Statistics, 46(5), 1961–1993.

Molina I, Rao JNK (2010). “Small Area Estimation of Poverty Indicators.” The Canadian Journal of Statistics, 38(3), 369–385. doi:10.1002/cjs.10051.

O’Sullivan, D., Unwin, D.J., (2010). Geographic Information Analysis, 2nd ed. John Wiley & Sons.

Parisi, F., Vangi, E., Francini, S., D’Amico, G., Chirici, G., Marchetti, M., Lombardi, F., Travaglini, D., Ravera, S., De Santis, E., Tognetti, R., 2023. S2 time series analysis for monitoring multi-taxon biodiversity in mountain beech forests. Front. Forests and Global Change 6. 10.3389/ffgc.2023.1020477.

Petrucci A, Salvati N (2004a) Small area estimation using spatial information, The Rathbun lake watershed case study. Working Paper no 2004/02, “G. Parenti” Department of Statistics, University of Florence

Petrucci A, Salvati N (2004b) Small area estimation considering spatially correlated errors: the unit level random effects model. Working Paper no 2004/10, “G. Parenti” Department of Statistics, University of Florence

Pinheiro J, Bates D, DebRoy S, Sarkar D, R Core Team (2018). nlme: Linear and Non-linear Mixed Effects Models. R package version 3.1-131.1, URL https://CRAN.R-project.org/package=nlme.

Rahlf, J., Breidenbach, J., Solberg, S., Næsset, E., & Astrup, R. (2014). Comparison of four types of 3D data for timber volume estimation. Remote Sensing of Environment, 155, 325–333.

Rao JNK, Molina I (2015). Small Area Estimation. 2.nd edition. Wiley Series in Survey Methodology, New Jersey: Wiley.

Roy, D. P., Boschetti, L., & Trigg, S. N. (2006). Remote sensing of fire severity: assessing the performance of the normalized burn ratio. IEEE Geoscience and remote sensing letters, 3(1), 112–116.

Rojas-Perilla, N., Pannier, S., Schmid, T., Tzavidis, N. Data-Driven Transformations in Small Area Estimation, Journal of the Royal Statistical Society Series A: Statistics in Society, Volume 183, Issue 1, January 2020, Pages 121–48, 10.1111/rssa.12488

Saarela S, Grafström A, Ståhl G, Kangas A, Holopainen M, Tuominen S, Nordkvist K, Hyyppä J (2015) Model-assisted estimation of growing stock volume using different combinations of LiDAR and Landsat data as auxiliary information. Remote Sens Environ 158:431–440

Saarela, S., Wästlund, A., Holmström, E., Mensah, A. A., Holm, S., Nilsson, M., … & Ståhl, G. (2020). Mapping aboveground biomass and its prediction uncertainty using LiDAR and field data, accounting for tree-level allometric and LiDAR model errors. Forest Ecosystems, 7(1), 43.

Shi, Y., Ke, G., Chen, Z., Zheng, S., & Liu, T.-Y. (2022). Quantized Training of Gradient Boosting Decision Trees. In S. Koyejo, S. Mohamed, A. Agarwal, D. Belgrave, K. Cho, & A. Oh (Eds.), Advances in Neural Information Processing Systems (Vol. 35, pp. 18822–18833). Curran Associates, Inc. https://proceedings.neurips.cc/paper_files/aper/2022/file/77911ed9e6e864ca1a3d165b2c3cb258-Paper-Conference.pdf.

Schmid, T., Tzavidis, N., Münnich, R., & Chambers, R. (2016). Outlier robust small-area estimation under spatial correlation. Scandinavian Journal of Statistics, 43, 806–826.

Sugasawa, S., Kubokawa, T. Small area estimation with mixed models: a review. Jpn J Stat Data Sci 3, 693–720 (2020). 10.1007/s42081-020-00076-x

Tabacchi, G., Di Cosmo, L., Gasparini, P., Morelli, S., 2011. Stima Del Volume E Della Fitomassa Delle Principali Specie Forestali Italiene, Equazioni Di Previsione, Tavole Del Volume E Tavole Della Fitomassa Arborea Epigea.

Tomppo, E., Olsson, H., Ståhl, G., Nilsson, M., Hagner, O., Katila, M., 2008. Combining national forest inventory field plots and remote sensing data for forest databases. Remote Sens. Environ. 112, 1982–1999. 10.1016/j.rse.2007.03.032.

Valavi, R., Elith, J., Lahoz-Monfort, J. J., & Guillera-Arroita, G. (2018). blockCV: An r package for generating spatially or environmentally separated folds for k-fold crossvalidation of species distribution models. Biorxiv, 357798.

Vangi, E., D’Amico, G., Francini, S., Giannetti, F., Lasserre, B., Marchetti, M., McRoberts, R.E., Chirici, G., 2021. The effect of forest mask quality in the wall-to-wall estimation of growing stock volume. Rem. Sens. 13, 1038. 10.3390/rs13051038

Ver Planck, N. R., Finley, A. O., Kershaw Jr, J. A., Weiskittel, A. R., & Kress, M. C. (2018). Hierarchical Bayesian models for small area estimation of forest variables using LiDAR. Remote Sensing of Environment, 204, 287–295.

White, J. C., Wulder, M. A., Hobart, G. W., Luther, J. E., Hermosilla, T., Griffiths, P., et al. (2014). Pixel-based image compositing for large-area dense time series

White, J. C., Arnett, J. T., Wulder, M. A., Tompalski, P., & Coops, N. C. (2015). Evaluating the impact of leaf-on and leaf-off airborne laser scanning data on the estimation of forest inventory attributes with the area-based approach. Canadian Journal of Forest Research, 45(11), 1498–1513.

White, J. C., Tompalski, P., Bater, C. W., Wulder, M. A., Fortin, M., Hennigar, C., … & White, R. (2025). Enhanced forest inventories in Canada: implementation, status, and research needs. Canadian Journal of Forest Research, 55, 1-37. applications and science. Can. J. Remote Sens. 40, 192–212. doi: 10.1080/07038992.2014.945827

Zhang, X., Shen, H., Huang, T., Wu, Y., Guo, B., Liu, Z., … & Ou, G. (2024). Improved random forest algorithms for increasing the accuracy of forest aboveground biomass estimation using Sentinel-2 imagery. Ecological Indicators, 159, 111752.

Zhang, X., Schaaf, C. B., Friedl, M. A., Strahler, A. H., Gao, F., & Hodges, J. C. (2002, June). MODIS tasseled cap transformation and its utility. In IEEE International Geoscience and Remote Sensing Symposium (Vol. 2, pp. 1063–1065). IEEE.

