## appendix of the main manuscript for "Small Area Estimation of Forest Volume Using Mixed Effects Random Forests and Multi-Source Remote Sensing Data"

In this section, the NER model's performance and other meaningful statistical metrics are reported, as well as the spatial scheme of the BLOOCV, the relationship between RMSE and sampling intensity and the partial dependence plot for the RF model.

 
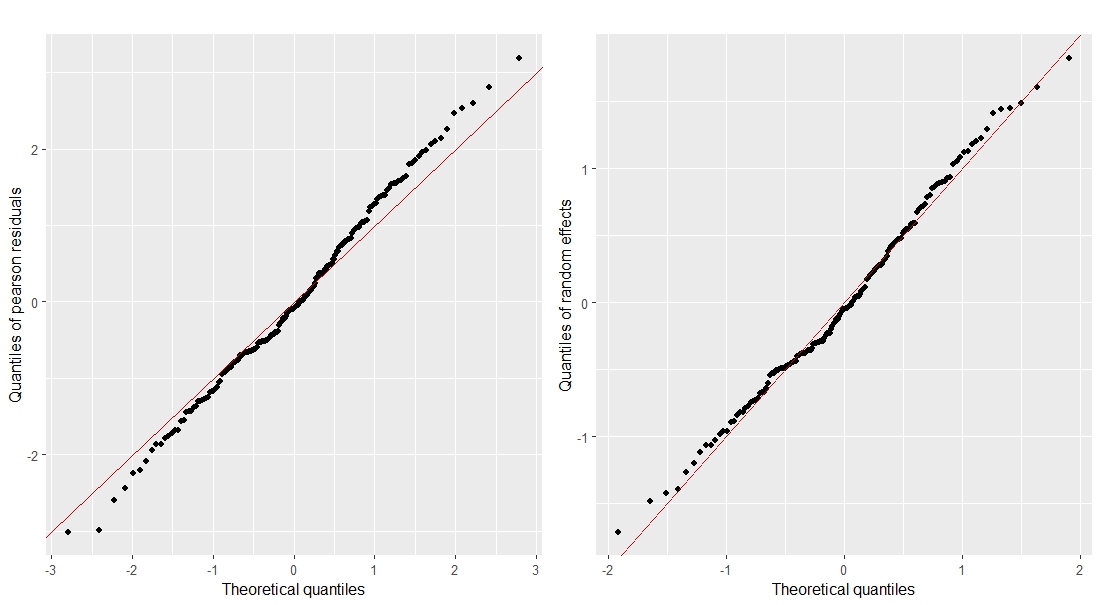

Figure S1. QQ-plot of fixed (left panel) and random (right panel) residuals.

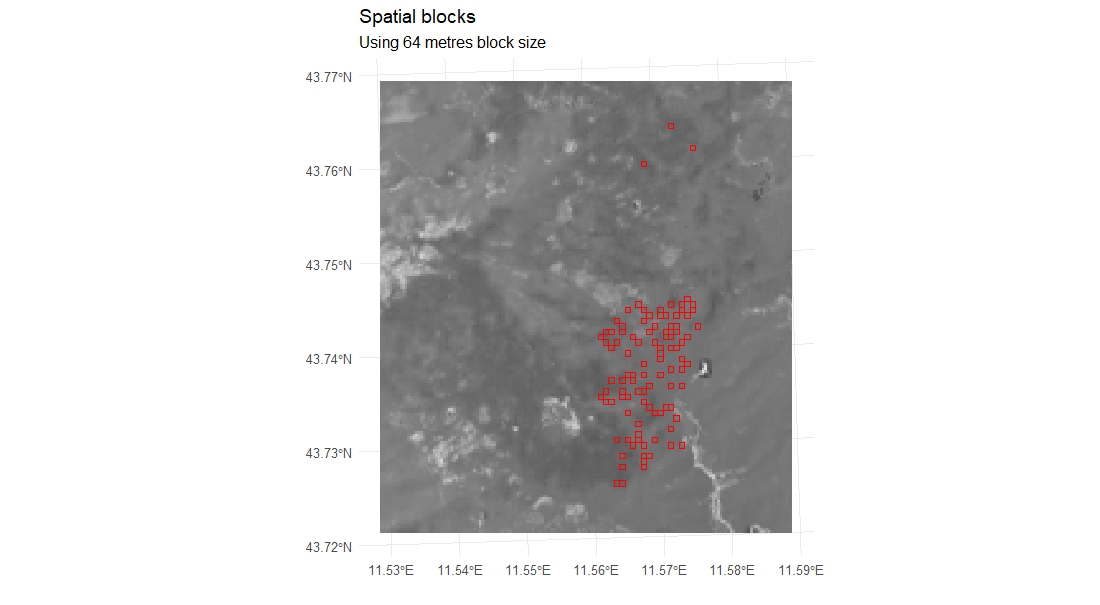

Figure S2. Spatial distribution of the 10 blocks generated for the cross-validation.

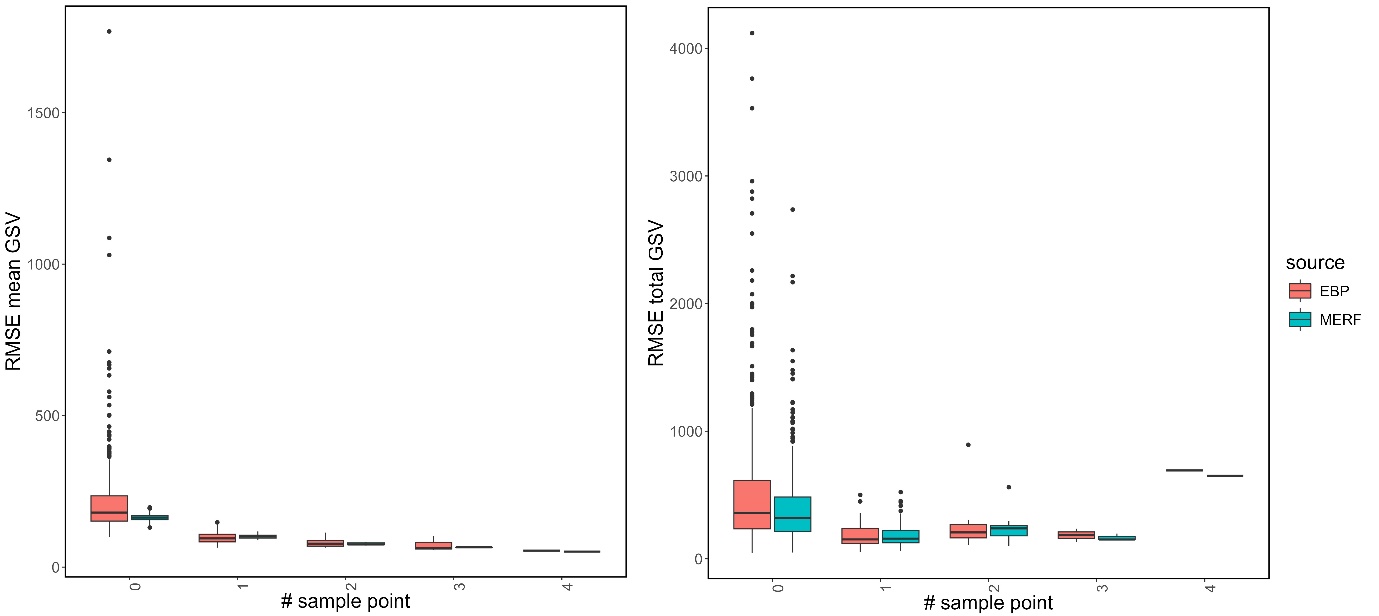

Figure S3. Distribution of RMSE for mean (left panel) and total (right panel) GSV at varying numbers of sampling plots within domains.

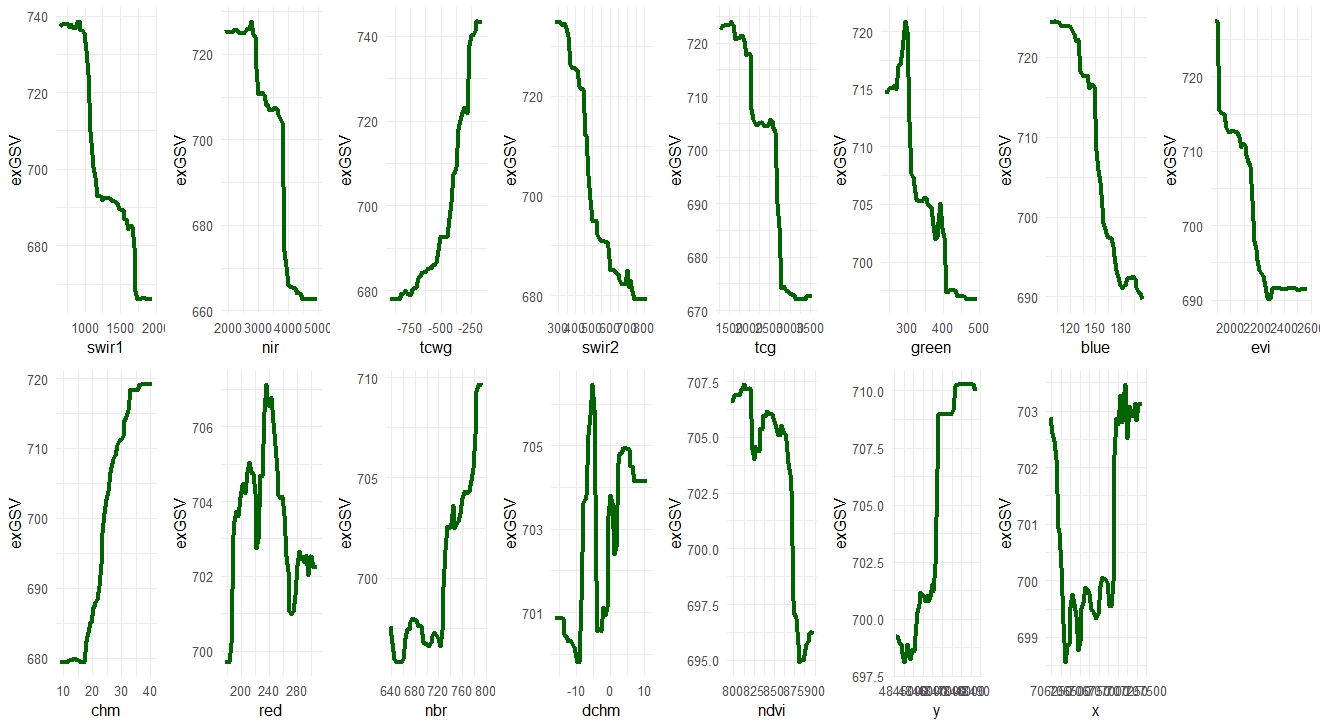

Figure S4. Partial dependence plot for variables ranked by their permutation importance
